# Influenza-specific effector memory B cells predict long-lived antibody responses to vaccination in humans

**DOI:** 10.1101/643973

**Authors:** Anoma Nellore, Esther Zumaquero, Christopher D. Scharer, Rodney G. King, Christopher M. Tipton, Christopher F. Fucile, Tian Mi, Betty Mousseau, John E. Bradley, Fen Zhou, Paul A. Goepfert, Jeremy M. Boss, Troy D. Randall, Ignacio Sanz, Alexander F. Rosenberg, Frances E. Lund

## Abstract

Seasonal influenza vaccination elicits hemagglutinin (HA)-specific CD27^+^ memory B cells (Bmem) that differ in expression of T-bet, BACH2 and TCF7. T-bet^hi^BACH2^lo^TCF7^lo^ Bmem are transcriptionally similar to effector-like memory cells while T-bet^lo^BACH2^+^TCF7^+^ Bmem exhibit stem-like central memory properties. T-bet^hi^ Bmem do not express plasma cell-specific transcription factors but do exhibit transcriptional, epigenetic, metabolic and functional changes that poise the cells for antibody production. Consistent with these changes, D7 HA^+^ T-bet^hi^ Bmem express intracellular immunoglobulin and T-bet^hi^ Bmem differentiate more rapidly into ASCs *in vitro.* The T-bet^hi^ Bmem response positively correlates with long-lived humoral immunity and clonotypes from T-bet^hi^ Bmem are represented in the early secondary ASC response to repeat vaccination, suggesting that this effector-like population can be used to predict vaccine durability and recall potential.

## Introduction

Following exposure to never before encountered antigens (Ag), the immune system initiates *de novo* responses with naive B and T cells. The progeny of the activated naive B cells can move into the germinal center (GC), engage with T_FH_ cells, undergo somatic hypermutation and affinity selection and, if appropriately signaled, will exit the GC and enter the long-lived memory B cell or plasma cell (PC) compartments (Elsner and Shlomchik, 2020). By contrast, when adults are exposed to Ags that are very closely related to Ags they have repeatedly encountered through natural infection or vaccination, the ensuing B cell response is dominated by pre-existing memory B cells (Andrews et al., 2015). In the case of virus-specific human memory B cells, experimental data (Turner et al., 2020) indicate that these cells can also enter the GC to produce new daughter cells that have the potential to become long-lived secondary PCs and memory cells. Memory B cells may also respond to vaccination or infection by proliferating outside of the GC and differentiate into short-lived plasmablasts (PB) (Lee et al., 2011). These terminally differentiated effectors, can reduce the duration of infection and decrease morbidity and mortality (Lee et al., 2011). The cues that direct memory B cells toward this early protective effector PB fate and the relationship between effector-like memory B cells and other durable GC-derived or extrafollicular memory B cell populations are not well understood (Laidlaw and Cyster, 2020).

Although the human memory B cell compartment is often subdivided into switched (IgD^neg^CD27^+^), unswitched (IgD^+^CD27^+^) and double negative (IgD^neg^CD27^neg^) memory B cell subsets, it is clear that considerable heterogeneity exists within these subpopulations (Sanz et al., 2019). In order to understand the molecular underpinnings of the memory B cell fate decisions made in response to infection or vaccination, analysis of Ag-specific memory B cells is required. Molecular studies of memory B cells have lagged behind those of memory T cells, in large part because Ag-specific memory B cells could only be identified retrospectively following *in vitro* differentiation of the memory B cells into ASCs (Boonyaratanakornkit and Taylor, 2019). With more widespread use of fluorochrome-labeled protein Ags, like recombinant influenza A virus hemagglutinin (HA) (Allie et al., 2019), it is now feasible to enumerate, isolate and characterize human Ag-specific B cells. Recent studies (Andrews et al., 2019; Ellebedy et al., 2016; Koutsakos et al., 2018; Lau et al., 2017) profiling the HA-specific B cell compartment after immunization with seasonal inactivated influenza vaccines (IIV) or following exposure to new HA Ags, like those derived from avian influenza strains, reveal significant heterogeneity within the memory B cell compartment. Indeed, vaccine-elicited B cells can be subdivided based on expression of the canonical memory B cell marker CD27 (Andrews et al., 2019) and on expression of activation markers, like CD71 (Ellebedy et al., 2016) and CD21 (Koutsakos et al., 2018; Lau et al., 2017), or inhibitory receptors like CD85j (Knox et al., 2017). Not surprisingly, the phenotypic variability observed within the HA-specific memory B cell compartment is accompanied by transcriptional heterogeneity between and within the different circulating HA-specific B cell subpopulations (Andrews et al., 2019; Lau et al., 2017).

The CD8 T cell memory compartment is also heterogeneous and can be divided into stem-like memory cells that retain considerable plasticity, effector-like memory T cells that can rapidly respond to Ag, and resident memory T cells that can provide local protection within tissues (Omilusik and Goldrath, 2019). The functional distinctions between these memory subsets are conferred by transcription factors (TF) that direct regulatory hubs. For example, BACH2 and TCF7 are critical for maintenance of the more stem-like properties of memory T cells (Pais Ferreira et al., 2020; Yao et al., 2021), while TFs like BATF and T-bet confer memory cells with effector potential (Kallies and Good-Jacobson, 2017; Kurachi et al., 2014). Many of the same fate-guiding TFs, including BACH2 (Hipp et al., 2017) and T-bet (Johnson et al., 2020; Zumaquero et al., 2019), are expressed by human B lineage cells. BACH2, which maintains memory B cell identity (Shinnakasu et al., 2016), must be repressed (Igarashi et al., 2014; Kometani et al., 2013) as B cells commit to becoming terminally differentiated antibody secreting cells (ASCs). By contrast, T-bet is required for memory B cell commitment to the ASC lineage as inducible deletion of T-bet specifically within the memory B cell compartment significantly impairs memory cell differentiation into ASCs following challenge infection with influenza (Stone et al., 2019). These data suggest that T-bet and BACH2 could play opposing roles in memory B cells, with BACH2 supporting the maintenance of the more stem-like memory cell pool and T-bet supporting effector potential within the memory compartment. Interestingly, T-bet expressing HA-specific memory B cells circulate in adults immunized with either seasonal Ag-drifted influenza strains (Lau et al., 2017) or given a prime/boost immunization with an avian H7 HA (Andrews et al., 2019). However, the T-bet-expressing human memory B cell subset has not been examined in detail because the markers typically used to subdivide memory B cells, like CD27 and CD21, do not resolve the T-bet expressing and non-expressing CD27^+^ memory B cells into discrete populations. For example, the HA-specific memory B cells found within the CD27^+^CD21^lo^ compartment on D14 post-IIV, which were characterized as pre-ASCs, includes both T-bet^+^ and T-bet^neg^ cells (Lau et al., 2017). Similarly, the classically defined CD27^+^IgD^neg^ memory HA-specific B cells that arise after prime/boost with H7 HA could be subdivided into three subsets – of which only two express T-bet (Andrews et al., 2019).

Here we interrogated the IIV-induced HA-specific T-bet^hi^ and T-bet^lo^ IgD^neg^CD27^+^ memory B cell compartments. We show that these two memory B cell subsets represent clonally, transcriptionally and epigenetically distinct populations. The T-bet^lo^ subset expresses stem-supportive TFs, like BACH2 and TCF7. By contrast, the T-bet^hi^ memory B cells have undergone T-bet associated epigenetic remodeling throughout the genome and specifically within the *BACH2* and *TCF7* loci. The T-bet^hi^ memory B cells, like ASCs, have largely repressed expression of *BACH2*, a known inhibitor of ASC differentiation (Igarashi et al., 2014). Unlike ASCs or pre-ASCs, T-bet^hi^ cells do not express ASC-specific TFs like *IRF4*, *BLIMP1* and *XBP1*. However, these cells do exhibit transcriptional changes consistent with the ASC effector fate including a switch toward mitochondrial fatty acid metabolism and oxidative phosphorylation, increased protein synthesis and catabolism and enhanced oxidative stress responses. These induced metabolic programs are associated with increased immunoglobulin (Ig) synthesis, activation of the mTORC1-dependent unfolded protein response (Gaudette et al., 2020) and rapid differentiation of the effector-like T-bet^hi^ memory B cell subset into ASCs. Importantly, the T-bet^hi^ effector memory B cell subset is a durable population that correlates with long-lived humoral immune responses following vaccination and can also rapidly contribute to secondary ASC responses in subsequent years.

## Results

### Influenza-specific T-bet^hi^ memory B cells are phenotypically and clonally distinct from T-bet^lo^ memory B cells

Our prior data, showing that antibody (Ab) responses by re-activated mouse memory B cells requires B cell intrinsic expression of T-bet (Stone et al., 2019), suggested to us that T-bet marks a population of memory B cells with effector potential. Although T-bet expressing B cells are rarely detected in circulation of healthy non-immunologically perturbed humans (Zumaquero et al., 2019) T-bet expressing B cells are observed in the blood of healthy donors (HD) following immunization with seasonal inactivated influenza virus (IIV) (Lau et al., 2017). To evaluate whether the vaccination-elicited T-bet expressing memory B cells exhibit effector-like properties, we used fluorochrome-conjugated HA H1 and H3 tetramers to enumerate circulating HA-specific B cells in HD immunized with seasonal IIV. Consistent with prior studies (Wrammert et al., 2008), we detected increased frequencies of circulating ASCs/PBs in the blood of day 7 (D7) vaccinated individuals (Fig. S1A-B). In addition, we identified IgD^neg^ H1^+^ B cells that could be divided (Fig. S1C) into T-bet^hi^ and T-bet^lo^ subpopulations (Fig. 1A) that were largely CD27^+^ (Fig. 1B), indicating that, at least in adults who have been repeatedly exposed to influenza through vaccination or infection, the early HA-specific B cell response to seasonal IIV is dominated by canonical CD27^+^IgD^neg^ memory cells. While both T-bet^hi^ and T-bet^lo^ H1 and H3-specific B cells were detected on D7 following IIV, we observed no correlation between the frequencies of the two populations in the vaccinated individuals (Fig. 1C-D). Moreover, the T-bet^hi^ and T-bet^lo^ populations differed in expression of other markers including FcRL5, CD21, CXCR5 and CXCR3 (Fig. 1B) as well as CD71, CCR7, CD85j and CD62L (Fig. S1D). Based on these markers, the CD27^+^IgD^neg^ T-bet^lo^ HA-specific memory B cells appeared most similar to resting memory (RM) cells while the CD27^+^IgD^neg^ T-bet^hi^ population appeared highly enriched in the CD71^+^CD85j^+^FcRL5^+^CD21^lo^CD62L^lo^CXCR3^+^CXCR5^neg^CCR7^neg^ activated memory 2 (AM2) subset that was identified following a booster vaccination with H7 HA Ag (Andrews et al., 2019).

**Figure 1.**
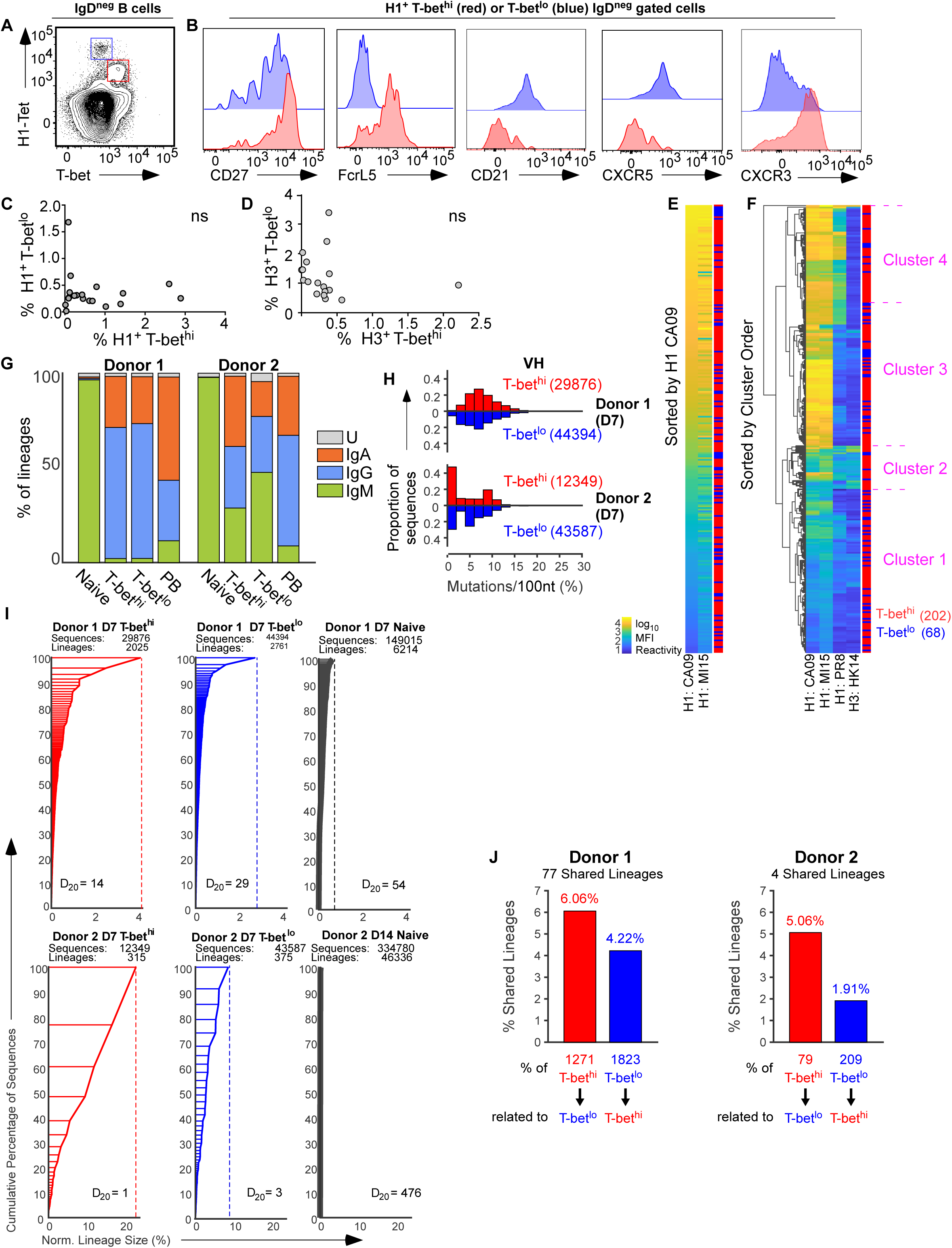
Seasonal influenza vaccination elicits distinct HA-specific memory B cell subsets. (**A-B**) Phenotype of B cell subsets (gating as in Fig. S1A, C) in peripheral blood of a representative HD on D7 post-immunization with seasonal IIV. T-bet expression (**A**) by H1 tetramer-binding B cells (H1^+^) IgD^neg^ gated B cells. Expression (**B**) of CD27, FcRL5, CD21, CXCR5 and CXCR3 (see Fig. S1D for additional markers) by the H1^+^ T-bet^hi^ (red) and H1^+^ T-bet^lo^ (blue) IgD^neg^ B cells. (**C-D**) Frequencies of T-bet^hi^ (X-axis) and T-bet^lo^ (Y-axis) memory cells within H1^+^ (**C**) or H3^+^ (**D**) IgD^neg^ B cell subsets measured in each individual (n=19) on D7 post-IIV (2015). ns = no significant correlation. (**E-F**) Single CA09-H1^+^ IgD^neg^ B cells from D7 post-IIV (2017) HD were index-sorted (see Fig. S1C for gating) and identified as either expressing high levels of T-bet (FcRL5^+^, n=202 cells, red) or low levels of T-bet (FcRL5^neg^, n=68 cells, blue). Recombinant IgG1 Abs (rAb) from sorted single cells were screened for binding (Table S1A) to H1 and H3 Ags arrayed on fluorochrome-labeled beads. Reactivity of each rAb for each H1 or H3 Ag represented as a heat map showing log10 mean fluorescence intensity (MFI) of binding after normalizing for binding to anti-IgG beads. Binding reactivities for each rAb to CA09-H1 and MI15-H1 (**E**) or CA09-H1, MI15-H1, PR8-H1 and HK14-H3 (**F**) were sorted by binding reactivity to CA09-H1 (**E**) or were hierarchically clustered using Euclidean distance and average linkage (**F**, Clusters 1-4). Distribution of the rMAbs derived from T-bet^hi^ cells (red) or T-bet^lo^ cells (blue) is indicated. See Table S1B for distribution of T-bet^hi^ and T-bet^lo^ cells within the clusters. (**G-J**) IGH variable region (IgH(V_H_)) BCR repertoire analysis of sort-purified (see Fig. S1A, C for gating) H3^+^ T- bet^hi^ and T-bet^lo^ IgD^neg^ B cells, IgD^+^ naïve B cells, and IgD^neg^CD38^hi^ PBs from peripheral blood of D7 HD (n=2) post-IIV (2016). (G) Isotype distribution in each subset (Table S1C). (H) V_H_ mutation load (Table S1D) in H3^+^ IgD^neg^ T-bet^hi^ and T-bet^lo^ cells. Mutations represented as frequency of mutations per 100 nucleotides (nt) in the V_H_ domain and in the CDR1+CDR2 or FR1+FR2+FR3 domains (Fig. S1E-F) of all Ig isotypes or from isotype-switched IgA and IgG B cells (Fig. S1G-I). Number of sequences in each population is indicated. (I) BCR repertoires shown as composition of lineages ordered by size (cumulative % of sequences, Y-axis) vs lineage size (shown as % of total number of sequences, X-axis). Diversity index 20 (D_20_) values, defined as the number of the largest size-ordered lineages that span the top 20% (cumulative) of sequences, are shown and total number of sequences and lineages for each subset are indicated. All lineages (Table S1E-F), including those with only one sequence (singletons), were identified in T-bet^hi^ and T-bet^lo^ populations. (J) Percentage of shared lineages between the T-bet^hi^ and T-bet^lo^ subsets excluding singleton lineages with the number of shared lineages (top) and total non-singleton lineages (bottom) indicated for each subset. Connections between lineages shown in Fig. S1J-K.

Since the vaccine-elicited CD27^+^T-bet^hi^ FcRL5^+^ AM2-like B cell subset has not been studied in detail, we interrogated this subset to determine whether these cells represent an effector memory population in humans. First, we isolated sort-purified single California 09 (CA09-H1^+^) IgD^neg^ B cells from a HD on D7 post-immunization and expressed the BCRs from these cells as recombinant Abs (rAbs). Next, we measured binding of the cloned rAbs to recombinant HA (rHA) Ags displayed on beads. As expected, rAbs generated from the cloned CA09-H1^+^ B cells bound CA09-H1 Ag arrayed beads and displayed a range of binding avidities (Fig. 1E, Table S1A). Recombinant Abs with high avidity for CA09-H1 Ag also reacted with high avidity to the closely related (97.4% identity) Michigan 2015 H1 Ag (MI15-H1) and rAbs that bound the CA09-H1 Ag with low avidity also reacted less well to the MI15-H1 Ag (Fig. 1E, Table S1A).

To measure the breadth of reactivity of the cloned CA09-H1-specific rAbs, we compared rAb binding to four rHA Ags, including CA09-H1, MI15-H1, PR8-H1 (81% identity to CA09-H1) and the heterosubtypic Hong Kong 2014 H3 Ag (HK14-H3). Hierarchical clustering of the reactivity data for each rAb (Fig. 1F and Table S1B) identified rAbs that bound with high (clusters 3, 4) or lower (clusters 1, 2) avidity to CA09-H1 and MI15-H1, rAbs that bound with high avidity to CA09-H1 and MI15-H1 and exhibited detectable binding to PR8-H1 (cluster 4) and broadly reactive rAbs that bound all 3 H1 Ags as well as the HK14-H3 Ag (cluster 2).

To determine whether the BCRs cloned from the T-bet^hi^ and T-bet^lo^ memory B cells exhibited distinct reactivity profiles, we utilized the index-sort information to quantitate FcRL5 expression levels (Fig. S1C) and assigned the cloned BCRs and rAbs to memory cells that expressed either high levels of T-bet (FcRL5^+^) or low levels of T-bet (FcRL5^neg^). rAbs with either high or low avidity for CA09-H1 were equally distributed within the vaccine-induced T-bet^hi^ and T-bet^lo^ H1^+^ IgD^neg^ B cell subsets (Fig. 1E). Moreover, we found no significant differences in the distribution of the rAbs cloned from T-bet^hi^ and T-bet^lo^ B cells within the different reactivity clusters (Fig. 1F, Table S1B). These data therefore indicate that high-avidity, low-avidity and broadly-reactive BCRs are present with similar distributions in both D7 Tbet^hi^ and T-bet^lo^ IgD^neg^ H1-specific B cell subsets.

Next, we purified RNA from sorted FcRL5^+^ (Tbet^hi^) and FcRL5^neg^ (T-bet^lo^) IgD^neg^ H3^+^ B cells from two D7 post-IIV HD (see Fig. S1C for sort strategy) and sequenced the rearranged Ig heavy chain variable (VH) region. Sorted IgD^+^CD27^neg^ naïve B cells and IgD^neg^CD27^hi^CD38^hi^ PBs from the same D7 donors (see Fig. S1A and S1C for sort strategy) were included as controls. Naïve B cells almost exclusively expressed IgM whereas most of the D7 PBs were isotype-switched to IgG or IgA (Fig. 1G, Table S1C). Consistent with a memory phenotype, switched IgG- and IgA-expressing B cells predominated in both T-bet^hi^ and T-bet^lo^ IgD^neg^ H3^+^ B cell compartments (Fig. 1G, Table S1C). While isotype usage varied between donors, the distribution of isotypes was similar in the T-bet^hi^ and T-bet^lo^ populations present within the same donor (Fig. 1G). Moreover, the entire VH region (Fig. 1H) as well as the CDR1+CD2 (Fig. S1E) and FR1+FR2+FR3 framework regions (Fig. S1F) of the expressed VH regions from the isotype switched and unswitched cells were mutated to a similar degree in the T-bet^hi^ and T-bet^lo^ IgD^neg^ H3^+^ B cell subsets (Table S1D). This was also true when the analysis was restricted to the isotype-switched B cells (Fig. S1G-I, Table S1D).

To assess whether the two populations included shared BCR clonotypes, we used repertoire analysis to identify lineages – defined as having an identical VH and J annotation, an identical HCDR3 length and HCDR3 nucleotide sequence similarity of >85% (Tipton et al., 2015) – within naïve, T-bet^hi^ and T-bet^lo^ IgD^neg^ H3^+^ B cells from the same donors. Naïve B cells exhibited high diversity (Fig. 1I) with many lineages. By contrast, the T-bet^hi^ and T-bet^lo^ subsets had fewer clonotypes and were less diverse than the bulk naïve B cells from the same individual (Fig. 1I), as indicated by a lower Diversity Index value (D_20_; (Tipton et al., 2015)), which represents the number of lineages in the top 20% when lineages are rank-ordered by size. Although large lineages were present in both T-bet^hi^ and T-bet^lo^ populations, the T-bet^hi^ H3^+^ population from each donor appeared to have larger lineages and a lower D_20_ score than the T-bet^lo^ H3^+^ cells from the same individual (Fig. 1I), suggesting that the T-bet^hi^ subset may be modestly more clonally restricted than the T-bet^lo^ subset. To address whether the lineages identified within the two memory B cell subsets were related, we determined the frequency of shared lineages between the two populations. In one donor we found that 6% of the T-bet^hi^ lineages were shared with lineages from the corresponding T-bet^lo^ subset and that 4% of the T-bet^lo^ clones were shared with clones from the corresponding T-bet^hi^ subset (Fig. 1J, Table S1E-F). Similar results were seen with the second donor (Fig. 1J, Table S1E-F). Interestingly, many of the larger clones found in the T-bet^hi^ and T-bet^lo^ populations did not overlap (Fig. S1J-K, Table S1E-F). Taken together, these results demonstrate that the vaccine-elicited D7 H3^+^ T-bet^hi^ and T-bet^lo^ IgD^neg^ B cells populations are bona fide memory cells that have undergone similar levels of isotype-switch, somatic mutation and clonal selection.

### HA-specific T-bet^hi^ and T-bet^lo^ IgD^neg^ B cells are transcriptionally and epigenetically distinct from pre-ASCs and ASCs

Prior reports revealed that human B cells expressing the integrin CD11c and lacking expression of CXCR5 and CD21 also co-express T-bet (Jenks et al., 2018; Wang et al., 2018; Zumaquero et al., 2019) This CD11c^+^CXCR5^neg^CD21^lo^ population is heterogenous and can be subdivided into at least three additional subsets, including IgD^+^CD27^neg^ activated naïve B cells, IgD^neg^CD27^neg^ CD11c^+^CXCR5^neg^ DN2 cells and IgD^neg^CD27^+^ cells (Sanz et al., 2019). Circulating DN2 cells, isolated from a subset of SLE patients, are reported to exhibit transcriptional and epigenetic properties of extrafollicular pre-ASCs (Jenks et al., 2018). Similarly CD27^+^IgD^neg^CD21^lo^ cells, found in circulation on Day 14 (D14) post-IIV, were described as pre-ASCs (Lau et al., 2017) and reported to express the ASC commitment TFs, IRF4 and BLIMP1. Given the phenotypic similarities between these known pre-ASC populations and the D7 vaccine-specific T-bet^hi^ IgD^neg^ cells, we considered the possibility that the T-bet^hi^ IgD^neg^ HA-specific cells were also pre-ASCs and derived from CD27^+^ memory compartment. To test this, we performed RNA-seq and ATAC-seq analyses on sort-purified naïve B cells, PBs, H1^+^ T-bet^hi^ (FcRL5^+^) and H1^+^ T-bet^lo^ (FcRL5^neg^) IgD^neg^ B cells (see gating in Fig. S1A and S1C) that were isolated from 5-6 HD on D7 post-IIV. Using a two-fold change (FC) in expression and a false discovery rate (FDR) cutoff of <0.05, we identified 762 differentially expressed genes (DEGs) and 1923 differentially accessible regions (DAR) in the chromatin between the T-bet^hi^ and T-bet^lo^ subsets (Table S2-3). As expected based on our sort strategy, *FCRL5* and *TBX21* were expressed at significantly higher levels in the T-bet^hi^ compared to the T-bet^lo^ IgD^neg^ H1^+^ B cells (Fig. S2A). To determine whether the transcriptome of the T-bet^hi^ H1^+^ B cells was enriched for genes that are expressed by the Lupus DN2 pre-ASCs (Jenks dataset (Jenks et al., 2018)) or vaccine-associated D14 CD21^lo^ pre-ASCs (Lau dataset (Lau et al., 2017)), we performed Gene Set Enrichment Analysis (GSEA). Consistent with our hypothesis, we observed (Fig. 2A) that DEGs, which are up in DN2 pre-ASCs relative to switched memory cells ((Jenks et al., 2018), Table S4) were significantly enriched in the T-bet^hi^ IgD^neg^ H1^+^ B cell transcriptome while DEGs that are downregulated in DN2 pre-ASCs (Table S4) were enriched in the T-bet^lo^ IgD^neg^ H1^+^ transcriptome. Similarly, DEGs that were up in the CD21^lo^ pre-ASCs compared to the CD21^hi^ B cells ((Lau et al., 2017), Table S4) were enriched in the T-bet^hi^ H1^+^ transcriptome and DEGs that were down in CD21^lo^ pre-ASCs (Table S4) were enriched in the T-bet^lo^ H1^+^ transcriptome (Fig. 2B). Despite this enrichment, <15% of the DEGs identified in the comparison between D7 H1^+^ IgD^neg^ T-bet^hi^ vs T-bet^lo^ B cells overlapped with the DEGs reported in the DN2 and CD21^lo^ pre-ASC datasets (Fig. 2C-D, Table S5A-B). These data suggest that the T-bet^hi^ IgD^neg^H1^+^ B cells exhibit substantial transcriptional differences from the previously described pre-ASC populations circulating in SLE patients and D14 IIV-vaccinated individuals.

**Figure 2.**
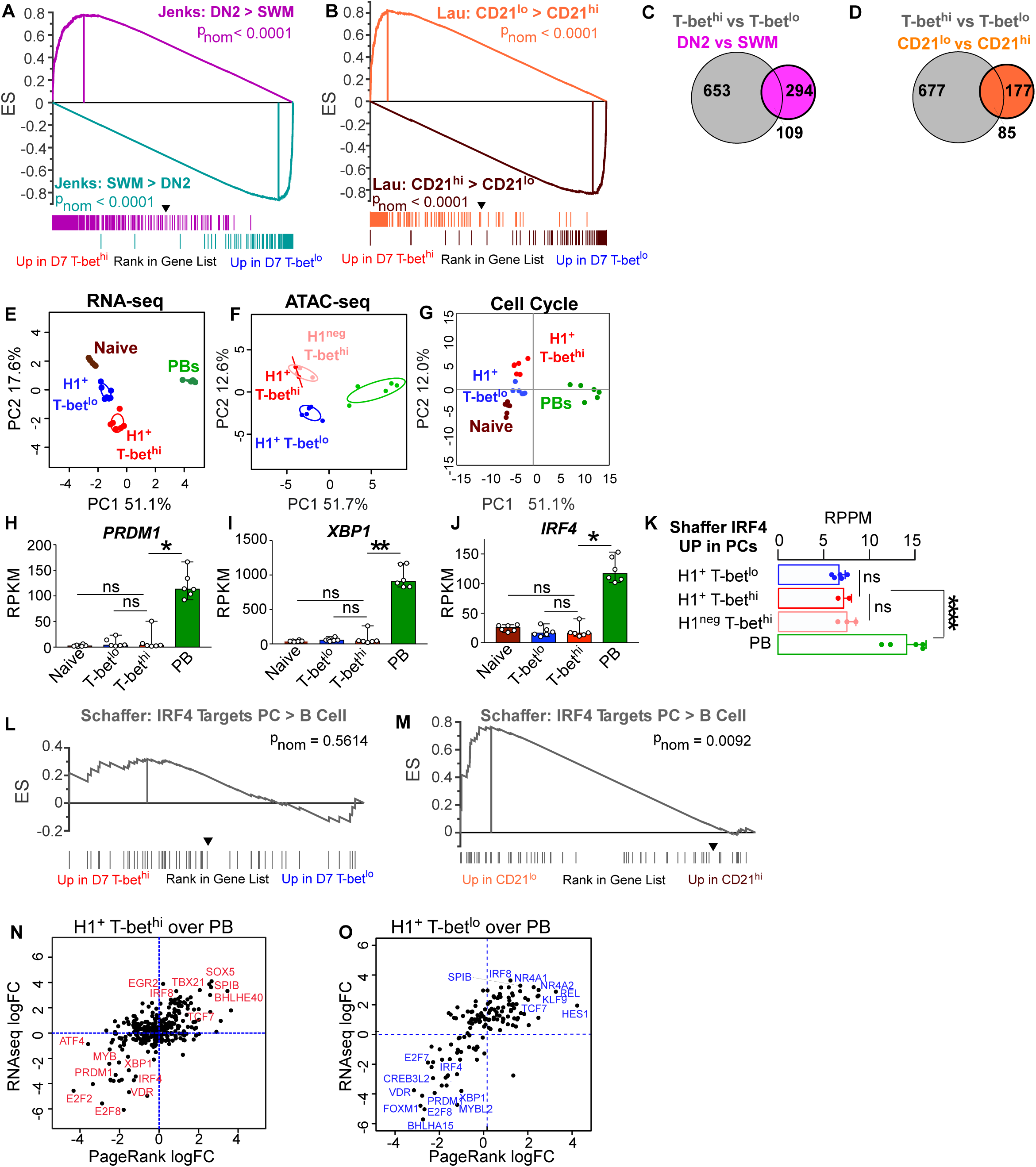
H1-specific T-bet^hi^ and T-bet^lo^ memory B cells are transcriptionally and epigenetically distinct from ASCs and pre-ASCs. (**A-O**) RNA-seq (Table S2) and ATAC-seq (Table S3) analyses performed on circulating PB (green), naïve B cells (brown), H1^+^ IgD^neg^ T-bet^lo^ (FCRL5^neg^, blue) cells, H1^+^ IgD^neg^ T-bet^hi^ B cells (FCRL5^+^, red), and bulk IgD^neg^ T-bet^hi^ (FCRL5^+^, pink) B cells that were sort-purified (Fig. S1A, C for sort-strategy) from HD (n=5-6) on D7 or D14 post-IIV. DEG and DAR defined as FDR<0.05 and FC >1 log(2) or <-1 log(2). T-bet and FcRL5 mRNA expression levels shown as Reads Per Kilobase per Million mapped reads (RPKM) shown in Fig. S2A. (**A-B**) GSEA comparing the RNA-seq ranked gene list from H1^+^ IgD^neg^ T-bet^hi^ and T-bet^lo^ memory B cells to (**A**) DEG that are over-expressed in T-bet-expressing IgD^neg^CD27^neg^ CD11c^+^CXCR5^neg^ B cells (DN2) compared to IgD^neg^CD27^+^ switched memory (SWM) B cells isolated from SLE patients (Table S4, (Jenks et al., 2018)) or (**B**) DEG identified as over-expressed in CD19^+^CD38^lo/med^ CD27^+^CD21^lo^ B cells relative to their CD21^hi^ counterparts on D14 post-IIV (Table S4, (Lau et al., 2017)). (**C-D**) Venn diagrams showing overlap between the 762 DEG identified in D7 H1^+^ IgD^neg^ T-bet^hi^ vs T-bet^lo^ memory B cells and (**C**) the 403 DEG identified between DN2 vs SWM B cells in SLE patients (Table S5A, (Jenks et al., 2018)) or (**D**) the 262 DEG identified between CD19^+^ CD38^lo/med^ CD27^+^CD21^lo^ vs CD21^hi^ B cells D14 post-IIV (Table S5B, (Lau et al., 2017)). Overlap analysis of D14 T-bet^hi^ and T-bet^lo^ memory B cell data sets (Table S5C-D) shown in Fig. S2H-I. (**E-F**) Principal component analysis (PCA) of the RNA-seq (**E**) and ATAC-seq (**F**) data sets from the indicated D7 B lineage subsets. D14 subsets shown in Fig. S2C-D. (**G**) PCA comparing expression of cell cycle genes (Table S4) in RNA-seq data from indicated D7 B cell subsets. Heat map in Fig. S2B. (**H**-**J**) RNA expression levels (RPKM) for *PRDM1* (**H**), *XBP1* (**I**) and *IRF4* (**J**) in indicated D7 B cell subsets. (**K**) Chromatin accessibility surrounding IRF4 regulated genes in human plasma cells (Shaffer et al., 2008) (Table S4) as assessed in ATAC-seq data derived from the indicated D7 (**K**) or D14 (Fig. S2E) B cell subsets. Data, reported as Reads per Peak per Million (RPPM), represent mean peak accessibility for all peaks mapping to genes directly regulated by IRF4 in plasma cells. (**L-M**) GSEA comparing the RNA-seq ranked gene list from D7 H1^+^ IgD^neg^ T-bet^hi^ vs Tbet^lo^ B cells **(L)** or the ranked gene list from CD19^+^CD27^+^CD38^lo/med^ CD21^lo^ vs CD21^hi^ B cells (Lau et al., 2017), **(M)** to known plasma cell-specific IRF4 targets (Shaffer et al., 2008). (**N-O**) TFs identified by PR (Table S6) as regulators of D7 PBs, D7 H1^+^ IgD^neg^ T-bet^hi^ and D7 H1^+^ IgD^neg^ T-bet^lo^ gene networks. Comparisons between the D7 T-bet^hi^ B cells over PB (**N**) or D7 T-bet^lo^ B cells over PBs (**O**) are shown with individual TFs indicated. D14 subsets (Table S6) in Fig S2F-G. Statistical analyses were performed using one-way ANOVA **(H-J**) and two tailed t-testing (**K**). *p< 0.05, **, p<0.01, *** p<0.001, **** p <0.0001 ns= non-significant. p_nom_ values for GSEA analyses are indicated.

Next, we performed Principal Component Analysis (PCA) comparing the transcriptomes (Fig. 2E, Table S2) and epigenomes (Fig. 2F, Table S3) of the D7 PBs to the D7 T-bet^hi^ and T-bet^lo^ IgD^neg^H1^+^ memory B cells. The T-bet^hi^ and T-bet^lo^ memory cells separated from the PBs on the PC1 axis when examining DEG or DAR. Cell cycle genes (Table S4), which were highly expressed by the proliferative D7 PBs but not by either T-bet^lo^ or T-bet^hi^ cells (Fig. 2G, Fig. S2B), were included in the DEGs between the PBs and memory cells. Likewise, the canonical ASC TFs *PRDM1*, *XBP1* and *IRF4* were highly expressed by the ASCs but were not expressed by naïve, T-bet^hi^ and T-bet^lo^ H1^+^ B cell subsets (Fig. 2H-J). Similarly, neither T-bet^hi^ nor T-bet^lo^ cells exhibited increased chromatin accessibility (Fig. 2K) surrounding genes that are expressed specifically in ASCs and directly regulated by IRF4 ((Shaffer et al., 2008), Table S4). Moreover, GSEA analysis (Fig. 2L) did not reveal any enrichment for expression of these IRF4-controlled ASC-specific genes in either the T-bet^hi^ or the T-bet^lo^ IgD^neg^ H1^+^ B cell transcriptomes. This was in contrast to DN2 pre-ASCs, which are reported to show significant enrichment for expression of IRF4-regulated ASC genes ((Jenks et al., 2018), and the D14 CD21^lo^ vaccine-elicited cells that were also enriched in expression of IRF4-controlled ASC-specific genes (Fig. 2M). We then used Page Rank (PR) analysis (Yu et al., 2017) to predict the key TFs that serve as regulatory hubs in each subset. As expected, the known ASC transcriptional regulators IRF4, PRDM1 and XBP1 were predicted to be modulators of the D7 PB gene expression program (Fig. 2N-O). However, these TFs were not predicted by PR to regulate the D7 T-bet^hi^ (Fig. 2N, Table S6) or T-bet^lo^ (Fig. 2O, Table S6) IgD^neg^ H1^+^ B cell transcriptional networks. Instead, TFs like IRF8 and SPIB, which are associated with B cell lineage identity (Schmidlin et al., 2008; Xu et al., 2015), were identified by PR as important regulators of gene expression in both T-bet^hi^ and T-bet^lo^ cells (Fig. 2N-O). Similar results were observed when we analyzed the transcriptomes and epigenomes of D14 T-bet^hi^ and T-bet^lo^ IgD^neg^H1^+^ B cells (Fig. S2C-G, Tables S2, S3, S6). Moreover, the overlap in DEGs between the previously described DN2 and CD21^lo^ pre-ASC subsets and the D14 vaccine-elicited T-bet^hi^ IgD^neg^H1^+^ memory B cells remained modest and did not include BLIMP1, IRF4 and XBP1 (Fig. S2H-I, Table S5C-D). Thus, our inability to identify a “pre-ASC” signature in the T-bet^hi^ memory population was not due to the timepoint chosen for analysis. Rather, despite phenotypic and transcriptional similarities between D7 T-bet^hi^ IgD^neg^ H1^+^ B cells and the previously described T-bet^hi^ DN2 and CD21^lo^ pre-ASC populations (Jenks et al., 2018; Lau et al., 2017), our data argued that neither population of memory B cells are transcriptionally programmed pre-ASCs.

### T-bet divides IgD^neg^ HA-specific B cells into distinct stable effector and stem-like memory populations

Circulating D7 T-bet^hi^ HA-specific IgD^neg^CD27^+^ B cells appeared to be *bona fide* memory cells rather than CD27^+^ memory-derived pre-ASCs. Given that T-bet expression by mouse memory B cells regulates the differentiation of these B cells into ASCs (Stone et al., 2019), we asked whether the vaccine-induced T-bet^hi^ memory B cells exhibited transcriptional or epigenetic changes that might support ASC development or function following reactivation. Interestingly, the D7 T-bet^lo^ IgD^neg^HA^+^ B cells expressed genes like *CCR7*, *BCL2* and *TCF7* (Fig. 3A, Table S2), which are more highly expressed by central memory T cells (Omilusik and Goldrath, 2019) while the D7 T-bet^hi^ IgD^neg^HA^+^ B cell subset expressed higher levels of genes such as *ZEB2*, *CXCR3*, and *TBX21* (Fig. 3A, Table S2), which are associated with T cell effector function (Omilusik and Goldrath, 2019). Furthermore, GSEA using curated gene lists of DEGs (Table S4) derived from published comparisons (Kaech et al., 2002; Luckey et al., 2006) between effector CD8 T cells and memory CD8 T cells revealed that the T-bet^hi^ memory B cells were significantly enriched for expression of genes that are more highly expressed by effector CD8 T cells compared to memory CD8 T cell genes (Fig. 3B). These results were further supported by PR analysis (Table S6) to identify TFs predicted to regulate the gene networks in D7 post-IIV T-bet^hi^ relative to T-bet^lo^ memory B cells. Indeed, “effector”-associated TFs (Kallies and Good-Jacobson, 2017; Kurachi et al., 2014), like T-bet and BATF, were predicted as regulators of the T-bet^hi^ memory B cell transcriptional gene network (Fig. 3C). By contrast, TFs associated with stem-like or central memory cells (Pais Ferreira et al., 2020; Zhou and Xue, 2012), like TCF7 and LEF1, were predicted as regulators of the T-bet^lo^ memory gene network (Fig. 3C).

**Figure 3.**
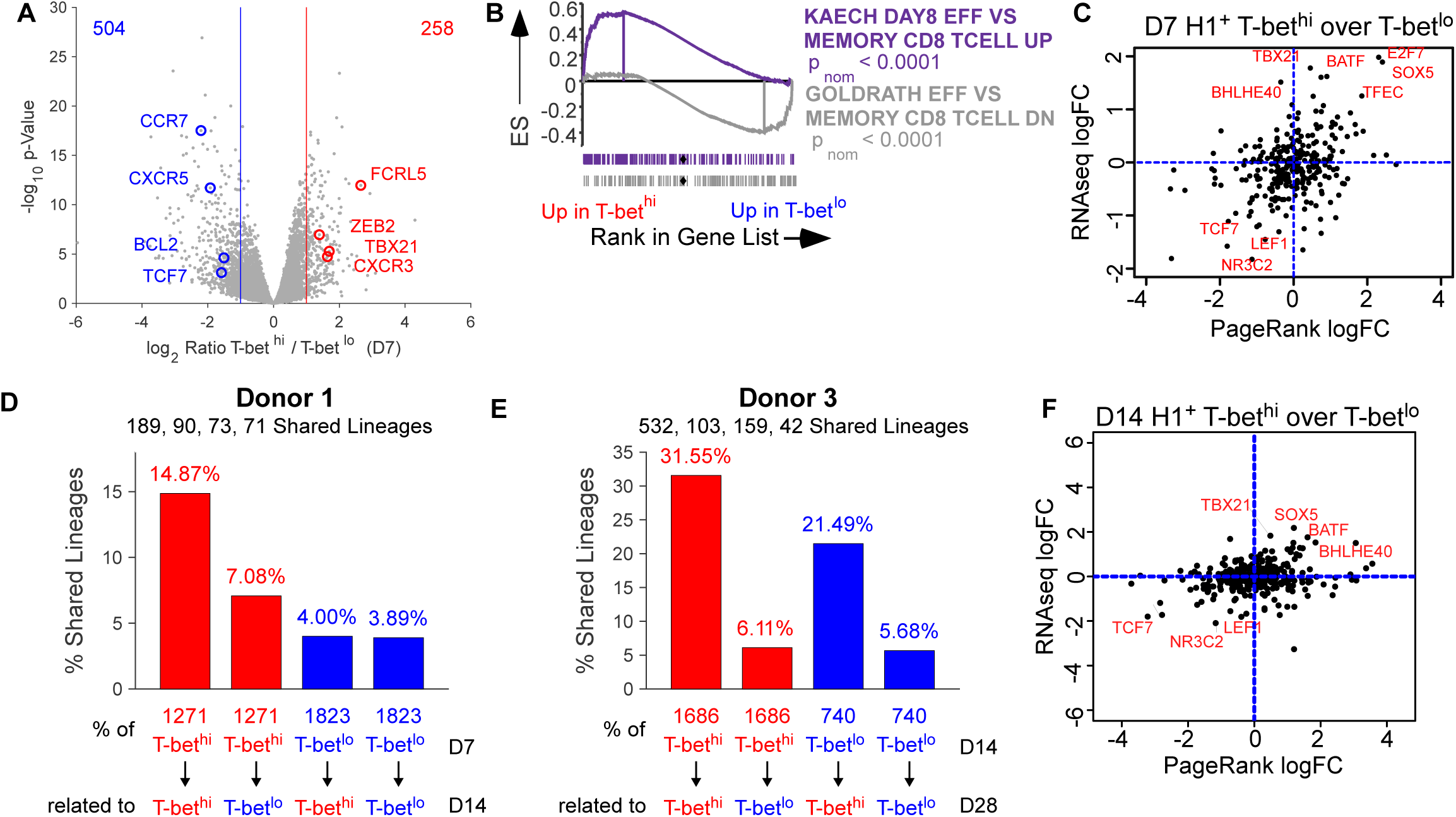
Division of H1-specific memory B cells into T-bet^hi^ effector-like and T-bet^lo^ stem-like subsets. **(A)** Volcano plot showing DEG upregulated in D7 H1^+^ IgD^neg^ T-bet^hi^ (red) and T-bet^lo^ (blue) memory B cells (Table S2) with callouts for specific DEG. **(B)** GSEA comparing the RNA-seq ranked gene list from D7 H1^+^ IgD^neg^ T-bet^hi^ and T-bet^lo^ B cells to DEG that are either upregulated (KAECH (Kaech et al., 2002)), Table S4) or down-regulated (GOLDRATH (Luckey et al., 2006)), Table S4) in effector vs memory CD8 T cells. **(C)** TFs identified by PR (Table S6) as regulators of D7 H1^+^ IgD^neg^ T-bet^hi^ and D7 H1^+^ IgD^neg^ T-bet^lo^ gene networks. Comparison between the T-bet^hi^ B cells over T-bet^lo^ B cells is shown with individual TFs indicated. (**D-E**) IgH(V_H_) BCR repertoire analysis to identify shared lineages between H1^+^ IgD^neg^ T-bet^hi^ and T-bet^lo^ B cells isolated from the same donor at sequential timepoints (Table S1E-F). Percentage of shared lineages shown with number of shared lineages between the various populations at different timepoints (top) and number of non-singleton lineages identified in each subset (bottom) indicated. **(F)** TFs identified by PR (Table S6) as regulators of D14 H1^+^ IgD^neg^ T-bet^hi^ and D14 H1^+^ IgD^neg^ T-bet^lo^ gene networks. Comparison between the T-bet^hi^ B cells over T-bet^lo^ B cells is shown and with individual TFs indicated.

One potential explanation for the transcriptional differences between the T-bet^hi^ and T-bet^lo^ HA^+^ B cells was that the T-bet^hi^ memory subset includes recently activated memory cells and the T-bet^lo^ memory B cell subset represents a more mature resting memory B cell population that accumulates with time. If so, then the clonotypes represented within the D7 T-bet^hi^ population might become enriched within the T-bet^lo^ subset as the cells transition into the more mature memory subset. To address this possibility, we determined the frequency of shared clones between circulating D7 T-bet^hi^ and T-bet^lo^ IgD^neg^ H3^+^ B cells and those present on D14 in the same individual. In contrast to the prediction, the lineages identified in the D7 T-bet^hi^ B cell compartment (Fig. 3D, Table S1E-F) were more represented in the D14 T-bet^hi^ population (14.9%) relative D14 T-bet^lo^ subset (7.1%). Similarly, comparison of the repertoires of D14 H1^+^ IgD^neg^ B cells and D28 H1^+^ IgD^neg^ B cells derived from a second donor (Fig. 3E, Table S1E-F) revealed many more connections between the D14 and D28 T-bet^hi^ subsets (31.6%) than between the D14 T-bet^hi^ and D28 T-bet^lo^ subsets (6.1%). Moreover, PR analysis (Table S6) using RNA-seq and ATAC-seq data derived from D14 T-bet^hi^ and T-bet^lo^ IgD^neg^HA^+^ B cells (Table S2-3) predicted LEF1 and TCF7 as regulators of the D14 T-bet^lo^ B cells and TBX21, BATF and BHLHE40 as regulators of the D14 T-bet^hi^ B cells (Fig. 3F). These data suggest that the T-bet^hi^ memory population is not a transient intermediate that transitions into the more mature T-bet^lo^ memory B cell compartment. Instead, the data suggest that seasonal influenza vaccination induces two clonally distinct, stable circulating memory B cell subsets that are endowed with different transcriptional and epigenetic programs.

### T-bet associated changes in effector memory B cells

Similar to T cells (Omilusik and Goldrath, 2019), vaccine-elicited memory B cells can be subdivided into effector cells that express high levels of T-bet and stem-like cells that express low levels of T-bet. T-bet can function as a transcriptional and epigenetic regulator (Oestreich and Weinmann, 2012). Given that PR analysis predicted T-bet as a regulator of 991 genes in the T-bet^hi^ memory B cell subset (Table S6), we hypothesized that T-bet likely contributes to the transcriptional and epigenetic landscape of the effector memory B cells. To assess this, we first enumerated the HOMER-defined T-bet binding motifs in DAR identified in the D7 ASCs, T-bet^hi^ and T-bet^lo^ memory B cells. We observed that chromatin accessibility surrounding T-bet binding motifs was significantly and differentially enriched in the T-bet^hi^ effector memory cells relative to the T-bet^lo^ memory cells and to the ASCs (Fig. 4A). Consistent with this, global analysis of the ENCODE T-bet ChIP-seq called peaks derived from GM12878 B lymphoblastoid cells (Consortium, 2012) revealed that known T-bet bound regions are significantly more accessible in the genome of the T-bet^hi^ memory B cell subset compared to the T-bet^lo^ cells or ASCs (Fig. 4B).

**Figure 4.**
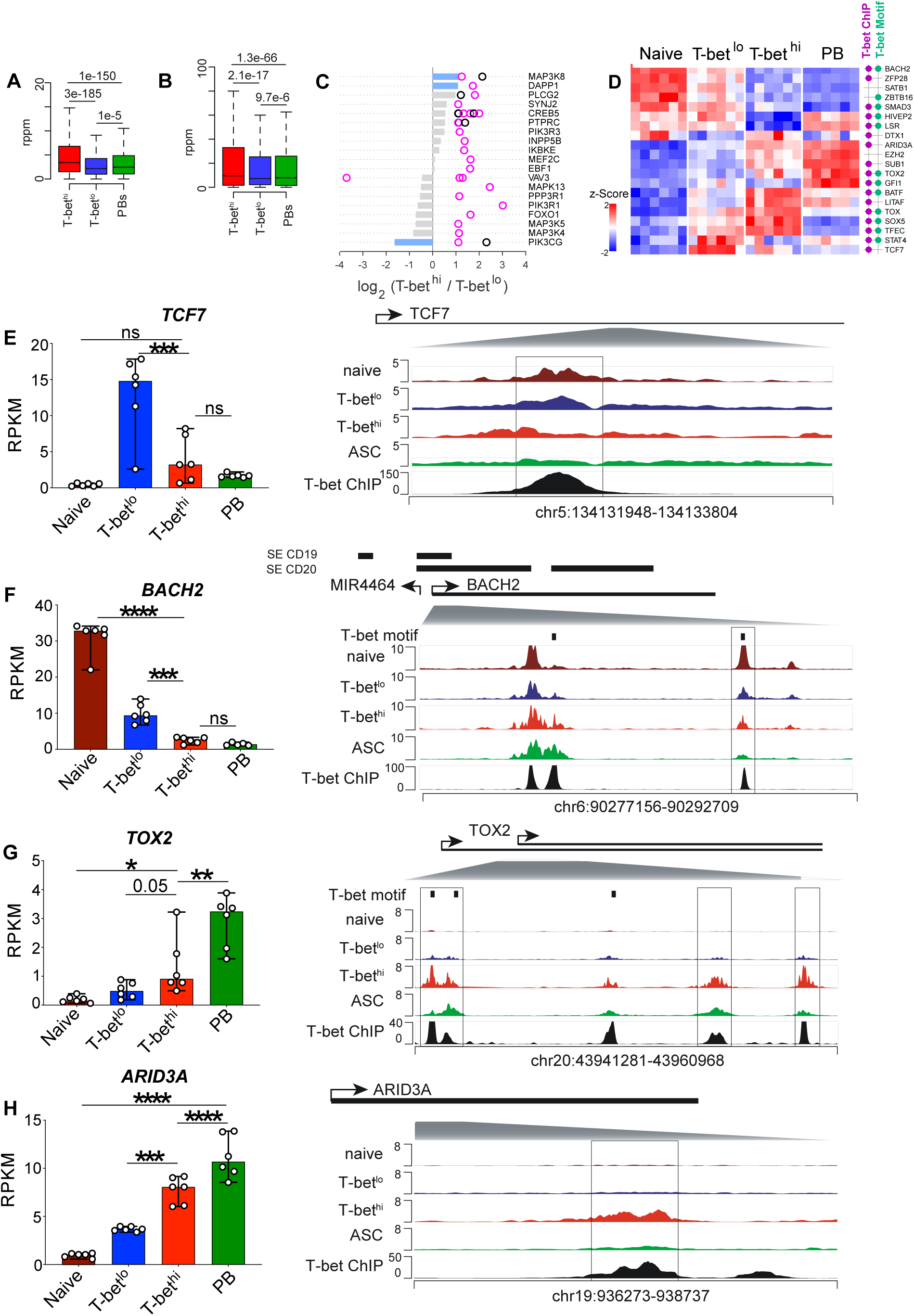
The effector memory B cell transcriptome and epigenome display T-bet associated changes. (**A-B**) Box plots of chromatin accessibility surrounding (within 50 bp) of defined T-bet motifs (**A**) and previously identified (Consortium, 2012) T-bet ChIP-seq peaks from GM12878 cells (**B**) in memory T-bet^hi^, T-bet^lo^ and PB populations. (C) A list of genes (Table S7) mapping to T-bet motif containing DAR that were increased (up) in T-bet^hi^ memory B cells were used in IPA (Table S8) and BCR signaling was identified as the top differentially regulated pathway between T-bet^hi^ and T-bet^lo^ memory B cells. Data are represented as the gene expression (bars) and chromatin accessibility (circles) in the 19/181 IPA-curated BCR signaling genes that are linked to increased chromatin accessibility regions containing T-bet binding motifs in the T-bet^hi^ memory subset. FC in gene expression shown in bars (blue bars significant DEG with at least 2-fold changes in expression up or down, grey bars not significant); DAR with T-bet binding motif (pink circle); DAR without T-bet binding motif (black circle). (D) Expression of 20 transcriptional regulators that were identified (Table S9) as DEG between T-bet^hi^ and T-bet^lo^ memory B cells and were assigned to at least one DAR between T-bet^hi^ and T-bet^lo^ cells. Expression levels in naïve, T-bet^hi^ memory, T-bet^lo^ memory and PB show as a z-score heat map. Genes assigned to DAR containing a T-bet binding motif (green dots) and/or to a DAR containing a ChIP-seq assigned (Consortium, 2012) T-bet binding site (purple dot) are indicated. (**E-H**) RNA-seq expression levels (in RPKM) and genome plots for four transcriptional regulators identified in (**D**) are shown. RNA expression levels for *TCF7* (**E**), *BACH2* (**F**), *TOX2* (**G**) and *ARID3A* (**H**) represented as bar graphs with the different B cell subsets and individual donors indicated. Genome plots aligned with T-bet binding sites as assessed by ChIP-seq of B lymphoblastoid cells (Consortium, 2012) and with ATAC-seq from resting naïve B cells (Scharer et al., 2019). Boxes indicate loci with significant DAR. Vertical black bars indicate consensus T-bet binding motifs, as predicted from HOMER. Super-enhancers identified in CD19^+^ and CD20^+^ human B cells (Hipp et al., 2017) are shown as horizontal black bars. Shaded gray triangles indicate location of DAR containing regions within each locus.

To determine whether these T-bet associated epigenetic changes were linked to particular effector pathways in the T-bet^hi^ memory B cells, we performed Ingenuity Pathway Analysis (IPA) on the 402 genes (Table S7) that were linked to at least one T-bet motif-containing DAR that was more accessible (increased) in the T-bet^hi^ memory B cells. IPA (Table S8) revealed BCR signaling as most differentially regulated pathway between T-bet^hi^ and T-bet^lo^ memory B cells. Although signaling pathways regulated by cytokines/cytokine receptors (e.g. IL-7, RANK) and growth factors (e.g. NGF, insulin) were also identified by IPA, each of these pathways was at least 1000-fold less enriched compared to BCR signaling in the T-bet^hi^ memory B cells. In fact, >10% of the IPA-defined BCR signaling genes were linked to at least one T-bet motif containing DAR that was more accessible in the T-bet^hi^ memory B cells (Fig. 4C). However, only three of these genes (*MAP3K8*, *DAPP1*, *PIK3CG*) were also differentially expressed by the T-bet^hi^ memory B cells (Fig. 4C). These data suggest that accessible T-bet motifs are associated with the epigenetic remodeling of BCR signaling genes in the T-bet^hi^ memory subset rather than with the differential expression of these genes.

To assess whether T-bet might more broadly contribute to effector memory B cell programming by regulating the expression of other TFs, we identified TFs that were differentially expressed between T-bet^hi^ and T-bet^lo^ memory B cells and were linked to a DAR in the T-bet^hi^ memory B cells (Table S9). Of the 20 TFs meeting these criteria, 75% were linked to a DAR associated with a T-bet binding motif and/or a T-bet ChIP-seq peak (Fig. 4D). Of particular interest, expression of a subset of these TFs changed progressively between T-bet^lo^ memory B cells, T-bet^hi^ memory B cells and ASCs (Fig. 4D). These included *TCF7*, which is reported to support stem-like potential in memory T cells (Pais Ferreira et al., 2020) and *BACH2*, which maintains B lineage identity (Shinnakasu et al., 2016) and prevents ASC development (Igarashi et al., 2014; Kometani et al., 2013). Consistent with a role in promoting stem-like potential, *TCF7* and *BACH2* were expressed at higher levels in the T-bet^lo^ memory B cells and exhibited progressively decreased gene expression and loss of chromatin accessibility within the DAR in T-bet^hi^ memory B cells and ASCs (Fig. 4E-F). By contrast, TFs like *TOX2* and *ARID3A,* which are most highly expressed in terminally differentiated ASCs showed progressively increased gene expression and enhanced chromatin accessibility within the associated DAR (Fig. 4G-H). Importantly, the DAR associated with each of these genes mapped to previously described T-bet ChIP-seq peaks (Fig. 4E-H,). These data are therefore consistent with the possibility that T-bet may facilitate epigenetic and transcriptional changes that bias the T-bet^hi^ memory B cells away from a stem-like program to the more effector-like program.

### T-bet^hi^ and T-bet^lo^ HA-specific memory B cells exhibit distinct metabolic and signaling profiles

Our data showed that the T-bet^hi^ effector memory subset expressed lower levels of *BACH2*, which maintains memory cells and prevents differentiation (Igarashi et al., 2014; Kometani et al., 2013) and higher levels of chromatin-modifying regulators, like *TOX2* and *EZH2*, that can support lymphocyte differentiation (Scharer et al., 2018; Xu et al., 2019). Based on these data, we predicted that the vaccine-induced T-bet^hi^Bach2^lo^ and T-bet^lo^Bach2^hi^ memory B cells subsets should exhibit distinct metabolic and functional profiles. To assess this in an unbiased way, we performed GSEA comparing the D7 T-bet^hi^ and T-bet^lo^ vaccine-specific memory B cell ranked gene list to gene sets in the curated Gene Ontology (GO) mSigDB gene set collection (Liberzon et al., 2011). Unexpectedly, none of the 3931 gene sets examined were significantly enriched in the T-bet^lo^ transcriptome. By contrast, we identified 185 gene sets in the GO collection that were enriched (FDR q< 0.01) in the T-bet^hi^ memory B cell transcriptome (Table S10). In each case, the leading-edge genes found in these enriched gene sets were expressed at higher levels in the T-bet^hi^ compared to T-bet^lo^ memory B cells (Table S10), suggesting that the pathways represented by these gene lists were likely to be activated in the T-bet^hi^ memory B cells. To evaluate this finding in more depth, we identified the leading-edge genes in each of the 185 gene sets and computed the Jaccard distance between each pair of leading-edge gene lists. After multidimensional scaling (MDS) followed by clustering, 9 clusters (Clusters A-I), each containing GO gene sets with overlapping leading-edge genes (Table S10, Fig. 5A), were identified. As expected, clusters that were in close proximity on the MDS plot contained related GO gene sets with overlapping leading-edge genes (Table S10). Using the clustering analysis as a guide, we selected 11 GO mSigDB gene lists ((Prototypic gene sets *i-xi*), Fig. S3A, Table S10) as representative of the gene sets found within the individual clusters. We then performed PCA analysis using the leading-edge genes identified by GSEA in the 11 prototypic GO gene lists. Not surprisingly, expression of these leading-edge differed substantially between D7 T-bet^hi^ and T-bet^lo^ HA^+^ memory B cells (Fig. 5B). Importantly, these transcriptional differences were maintained for at least 14 days (Fig. 5B), indicating that transcriptional differences between the two populations of memory cells revealed by GSEA were intrinsic and durable.

**Figure 5.**
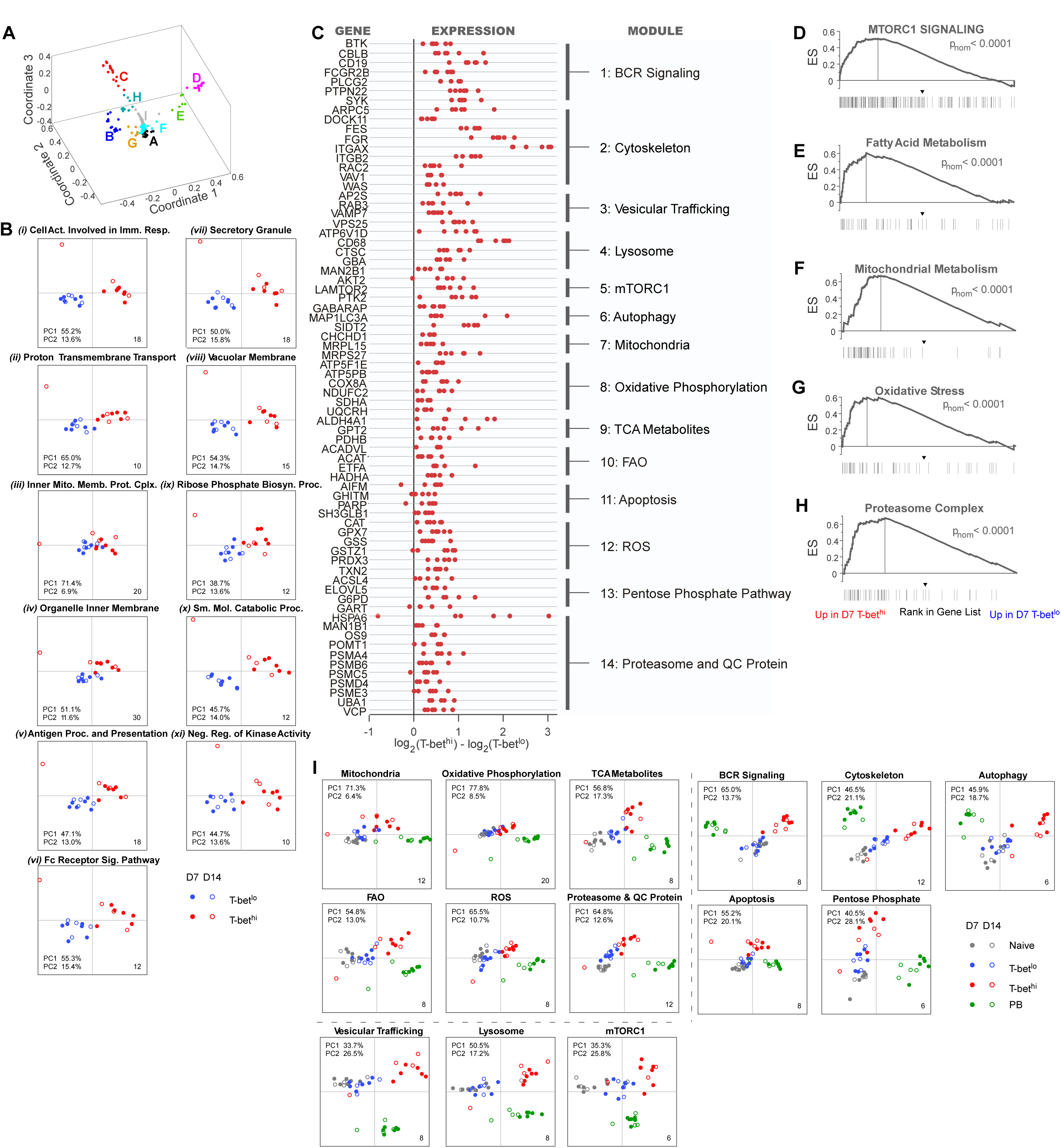
Metabolic gene expression differs between T-bet^hi^ effector and T-bet^lo^ stem-like memory B cells. **(A)** GSEA comparing the RNA-seq ranked gene list from D7 H1^+^ IgD^neg^ T-bet^hi^ and T-bet^lo^ B cells to 3931 (min gene set size = 15, max gene set size = 500) GO gene sets (mSigDB v.7) was performed. GO gene lists (n = 185) significantly enriched (FDR q-value < 0.01,) for expression in T-bet^hi^ relative to the T-bet^lo^ memory B cells were analyzed by multidimensional scaling and clustered (9 clusters (A-I), Table S10; see methods). (B) Eleven prototypic GO gene lists (*i-xi*) representing the 9 GO gene set clusters were selected (Table S10, Fig. S3A). Leading-edge genes from each of the 11 GO gene lists (Table S10) were assessed by PCA comparing H1^+^ IgD^neg^ T-bet^hi^ (red) and T-bet^lo^ (blue) memory B cells on D7 (closed circle) and D14 (open circle) post-IIV. (C) Leading-edge genes from the 11 prototypic GO gene lists were curated into 14 signaling and metabolic modules (Table S11). Expression of representative genes from each module by D7 H1^+^ IgD^neg^ T-bet^hi^ and T-bet^lo^ B cells is shown. All genes in each module provided in Fig. S3B-O). Red dots indicate individual RNA-seq samples with FDR q<0.05. (**D-J**) GSEA comparing the RNA-seq ranked gene list from D7 H1^+^ IgD^neg^ T-bet^hi^ and T-bet^lo^ B cells to curated gene lists (Table S4) for mTORC1 signaling **(D)**, fatty acid metabolism **(E)**, mitochondrial metabolism **(F)**, oxidative stress **(G)** and proteasome complex **(H)**. p_nom_ values are indicated. **(I)** Genes from 14 modules (name above each plot) assessed by PCA comparing naïve B (grey), PB (green), H1^+^ IgD^neg^ T-bet^hi^ (red) and H1^+^ IgD^neg^ T-bet^lo^ (blue) from memory B cells on D7 (closed circle) and D14 (open circle) post-IIV.

Consistent with our hypothesis, many of the transcriptional differences between the T-bet^hi^ and T-bet^lo^ memory B cells were associated with GO terms linked to immune cell function and metabolism (Fig. 5B, Fig. S3A). To better characterize these differences, we curated the functional annotations of the leading-edge genes found in the 11 prototypic GO gene lists and grouped them into 14 signaling and metabolic genes modules that were upregulated in the T-bet^hi^ memory B cell subset ((module 1-14) Fig. 5C, Table S11). In agreement with our earlier IPA analysis (Table S8), Module 1 included genes encoding proteins associated with BCR signaling (Fig. 5C, Fig. S3B, Table S11). Module 2 included genes associated with cytoskeleton assembly and signaling (Fig. 5C, Fig. S3C, Table S11), like integrins (e.g. CD11c, *ITGAX* and *ITGB2*), small G proteins (e.g. *RAC2*), guanine nucleotide exchange factors (e.g. *DOCK11* and *VAV1*), protein kinases (e.g. *FGR* and *FES*) and members of the WASP/ARP2/3 complex (*WAS* and *APRC5*). Module 3 included genes like *AP2S1*, *VPS25*, *RAB34* and *VAMP7* that are involved trafficking, sorting and fusion of cargo-containing vesicles to endosomes and lysosomes (Fig. 5C, Fig. S3D, Table S11). Module 4 included genes involved in lysosome function (Fig. 5C, Fig.S3E, Table S11), like scavenger receptors (e.g. *CD68*), subunits of the lysosome ATPase (e.g. *ATP6V1D*) and enzymes required for peptide (e.g. *CTSC*), ceramide (e.g. *GBA*) and carbohydrate (e.g. *MAN2B1*) degradation. These data suggested that macromolecule trafficking and endo-lysosomal activity are increased in T-bet^hi^ memory cells.

The endosomal-lysosomal compartment serves as the site for protein sorting, degradation and recycling. In addition, the lysosome supports the mTORC1 complex, which functions as a cell sensor for nutrients (Saxton and Sabatini, 2017). Consistent with published data showing that the mTORC1 complex is preferentially activated in effector memory relative to central memory CD8 T cells (Buck et al., 2017), we observed increased expression of genes associated with mTORC1 (Module 5, Fig. 5C, Fig. S3F, Table S11) in the T-bet^hi^ “effector-like” memory B cells. These included genes encoding activators of mTORC1 (e.g. *AKT2, PTK2*) and genes encoding subunits of the ragulator complex (e.g. *LAMTOR2*), which recruits mTORC1 to the lysosome. In accordance with the role of mTORC1 regulating the balance between protein synthesis and autophagy (Saxton and Sabatini, 2017), genes that catalyze the transfer of amino acids to tRNA molecules (tRNA synthetase genes, e.g. *QARS*, *WARS* Table S10 Cluster H) were also increased in the T-bet^hi^ memory B cells. Likewise, genes involved in the formation, maturation and function of autophagosomes (Module 6, Fig. 5C, Fig. S3G, Table S11) like *GABARAP*, *MAP1LC3A*, *SIDT2* were increased in T-bet^hi^ memory B cells. These data suggest that T-bet^hi^ memory B cells have enhanced capacity to both synthesize and degrade proteins.

Protein synthesis catalyzed by tRNA synthetases requires ATP. Module 7 included genes that regulate the activity and function of the ATP-generating mitochondria (Fig. 5C, Fig. S3H, Table S11), like the large and small mitochondrial ribosome subunits (e.g. *MRPL15* and *MRPS27*) and proteins involved in mitochondrial biogenesis and structure (e.g. *CHCHD1*). Module 8 included genes associated with the mitochondrial electron transport chain (ETC) and oxidative phosphorylation (Fig. 5C, Fig. S3I, Table S11) like members of the mitochondrial ATP synthase complex (e.g. *ATP5F1* and *ATP5E*), the mitochondrial inner membrane Complex I-III (e.g. *NDUFC2*, *SDHA* and *UQCRH*) and Complex IV Cytochrome c oxidase (e.g. *COX8A*). Consistent with this, Module 9 included genes controlling the production or transport of metabolites that feed the tricarboxcyclic acid (TCA) cycle (Fig. 5C, Fig. S3J, Table S11) like *ALDH4A*, *PDHB* and *GPT2*. Module 10 included genes encoding enzymes required for mitochondrial fatty acid β-oxidation (FAO, Fig. 5C, Fig. S3K, Table S11) like *ACADVL*, *ACAT1*, *ETFA* and *HADHA*. Since FAO produces metabolites like acetyl-CoA, NADH and FADH, which drive the TCA cycle and are used as redox partners in the ETC, the data suggested that the T-bet^hi^ memory B cells may preferentially utilize oxidative phosphorylation to generate ATP.

Fatty acid catabolism and ATP production during oxidative phosphorylation results in generation of reactive oxygen species (ROS) that regulate signaling, differentiation and cell death (Bertolotti et al., 2012). Consistent with their enhanced oxidative phosphorylation signature (Modules 7-10), T-bet^hi^ memory B cells also expressed increased levels of genes associated with caspase-dependent and independent apoptosis (Module 11, Fig. 5C, Fig. S3L, Table S11) like *AIFM1*, *PARP1* and *GHITM*. Conversely, T-bet^hi^ memory B cells expressed higher levels of genes associated with detoxifying free radicals (Module 12, Fig. 5C, Fig. S3M, Table S11), like peroxiredoxins (e.g. *PRDX3*), thioredoxins (e.g. *TXN2*), catalase (e.g. *CAT*) and enzymes in the pentose phosphate pathway (PPP)-driven glutathione redox reactions (e.g. *GSTZ1)*. Other genes associated with the PPP (Module 13, Fig. 5C, Fig. S3N, Table S11), including glucose-6-phosphate dehydrogenase (*G6PD*), which catalyzes the rate limiting step in shunting glucose from glycolysis to PPP and enzymes involved in PPP-dependent purine (e.g. *GART*) and FA biosynthesis (e.g. *ELOVL5* and *ACSL4*) were also upregulated in T-bet^hi^ memory cells. Finally, Module 14 (Fig. 5C, Fig. S3O, Table S11) included genes involved in endoplasmic reticulum (ER) protein folding and transport (e.g. *HSPA6*) and enzymes that regulate N-linked glycosylation of newly translated proteins in the ER (e.g. *DOST*). However, T-bet^hi^ memory B cells also expressed increased levels of *UBA1* (Fig. 5C), which is the activating E1 ubiquitin enzyme found in the ER associated degradation complex (ERAD, (Oikonomou and Hendershot, 2020)), and *VCP* (Fig. 5C), which is the cytoplasmic AAA+ ATPase that facilitates translocation of misfolded proteins from the ER back into the cytoplasm where these proteins are targeted to the proteasome and degraded. Moreover, T-bet^hi^ memory B cells expressed higher levels of genes encoding ER proteins (e.g. *MAN1B1*, Fig. 5C) that demannosylate misfolded glycoproteins and recognize (*OS9*, Fig. 5C) and target these mannose-trimmed proteins to the ERAD complex. Finally, proteasome and immunoproteasome subunits (e.g. *PSMA4*, *PSMB6*, *PSMC5*, *PSMD4, PSME3*) that are required for proteasome-mediated protein degradation were upregulated in the T-bet^hi^ memory B cells (Fig. 5C). Collectively, these data suggest that T-bet^hi^ memory B cells are transcriptionally programmed to promote the synthesis and degradation of proteins and other macromolecules and have engaged multiple mechanisms to combat the oxidative stress associated with these processes.

Our curated bioinformatic analysis showed that the H1-specific T-bet^hi^ and T-bet^lo^ memory B cells differed in their expression of genes associated with numerous metabolic processes, including those linked to mTORC1 activity (Module 5-6), mitochondrial function (Modules 7-9), FAO (Module 10), PPP and ROS detoxification (Modules 12-13) and proteasome activity (Module 14). To validate these results, we performed additional GSEA using independently curated gene lists (Table S4) for these metabolic pathways. Importantly, and consistent with our conclusions, the transcriptome of the T-bet^hi^ memory B cells was highly enriched (p_nom_ <0.0001) for expression of genes associated with mTORC1 signaling (Fig. 5D), fatty acid metabolism (Fig. 5E), mitochondrial metabolism (Fig. 5F), oxidative stress (Fig. 5G) and the proteasome (Fig. 5H).

To assess whether these signaling and metabolic transcription modules were stable in the T-bet^hi^ memory B cells, we performed PCA with the gene lists (Table S11) from the 14 modules using transcriptome datasets from D7 and D14 T-bet^hi^ and T-bet^lo^ memory B cells, naïve cells and PBs. The differences between expression of genes in the metabolic and signaling transcription modules were durable and persisted over at least 14 days in the vaccine-induced HA-specific T-bet^hi^ and T-bet^lo^ memory B cell populations (Fig. 5I). Moreover, the T-bet^lo^ memory B cells were transcriptionally more similar to the naïve B cells in expression of the genes within the 14 modules (Fig. 5I). By contrast, the T-bet^hi^ memory B cells did not precisely co-localize with either the naïve/T-bet^lo^ cluster or the ASCs (Fig. 5I). In some cases, the T-bet^hi^ memory cells co-segregated with the ASCs on either the PC1 axis (modules for vesicular trafficking, lysosomal activity, and mTORC1) or the PC2 axis (modules for BCR signaling, cytoskeleton and autophagy modules) (Fig. 5I). In other cases (modules for mitochondria, oxidative phosphorylation, TCA, FAO, ROS and proteasome), we observed a gradient in gene expression along the PC1 axis that proceeded from naïve B cells, followed closely by T-bet^lo^ memory B cells, to T-bet^hi^ memory B cells and finally to ASCs (Fig. 5I). This gradient of gene expression was reminiscent of that seen for expression of *TCF7*, *BACH2*, *ARID3A* and *TOX2*. Importantly, in each module the T-bet^lo^ memory cells clustered more closely to the naïve B cells while the T-bet^hi^ cells exhibited transcriptional characteristics that were more similar but not identical to the PBs. These data therefore indicate that the metabolic transcriptional profiles of vaccine-induced T-bet^hi^ and T-bet^lo^ memory B cells are distinct, stable and intrinsic to the two populations. Moreover, while vaccine-induced H1-specific T-bet^hi^ CD27^+^IgD^neg^ memory B cells are not transcriptionally programmed pre-ASCs, this population of memory B cells does exhibit increased expression of genes controlling metabolic processes associated with the high energy demands required for Ab synthesis and secretion by ASCs.

### T-bet^hi^ memory B cells are metabolically poised to rapidly differentiate upon reactivation

Transcriptome analysis of the vaccine-specific T-bet^hi^ and T-bet^lo^ memory B cells indicated that these cells differed in expression of genes involved in multiple metabolic processes. We therefore predicted that these transcriptional differences would translate into metabolic changes between the two memory B cell subsets. Because the low numbers of the Ag-specific memory B cell subsets in blood precluded most functional studies, we first evaluated the T-bet^hi^ and T-bet^lo^ CD27^+^IgD^neg^ classical memory B cells that are present at low levels in adult human lymphoid tissues (Li et al., 2016). Consistent with the prior data (Li et al., 2016), the CD27^+^IgD^neg^ T-bet^hi^ memory cells from human tonsil were uniformly FcRL5^+^ (Fig. S4A-C). We therefore used this marker to subdivide the tissue-residing memory B cells into T-bet expressing (FcRL5^+^) or non-expressing (FcRL5^neg^) populations. To assess whether the tissue-derived memory CD27^+^ T-bet^hi^ and T-bet^lo^ cells exhibited differences in their metabolic profiles we stained the cells with either H_2_DCF to measure intracellular ROS levels (Fig. 6A, Fig. S4D) or with the mitochondrial matrix dye mitotracker green (Fig. 6B, Fig. S4E), which detects oxidative and nitrosative stress within cells and is often used as a surrogate measure of mitochondrial mass (Doherty and Perl, 2017). In both cases (Fig. 6A-B), the T-bet^hi^ memory B cells stained more intensely, thereby suggesting that metabolic activity differs between tissue-derived T-bet^hi^ and T-bet^lo^ memory B cells. Consistent with this conclusion, the T-bet^hi^ memory B cells exhibited changes in fatty acid metabolism as these cells stained more intensely with BODIPY 510 (Fig. 6C, Fig. S4F), which measures incorporation of fatty acids into phosphatidyl choline and the subsequent breakdown of these metabolites into di- and tri-glycerides. Importantly, the T-bet^hi^ memory cell subset also exhibited increased basal phosphorylation of S6 kinase (Fig. 6D, Fig. S4G), a direct downstream target of the metabolic sensor mTORC1. These data reveal that the tissue residing memory T-bet^hi^ B cell subset displays functional attributes that are in alignment with the predicted metabolic profiles identified in the vaccine-specific circulating T-bet^hi^ memory B cell population.

**Figure 6.**
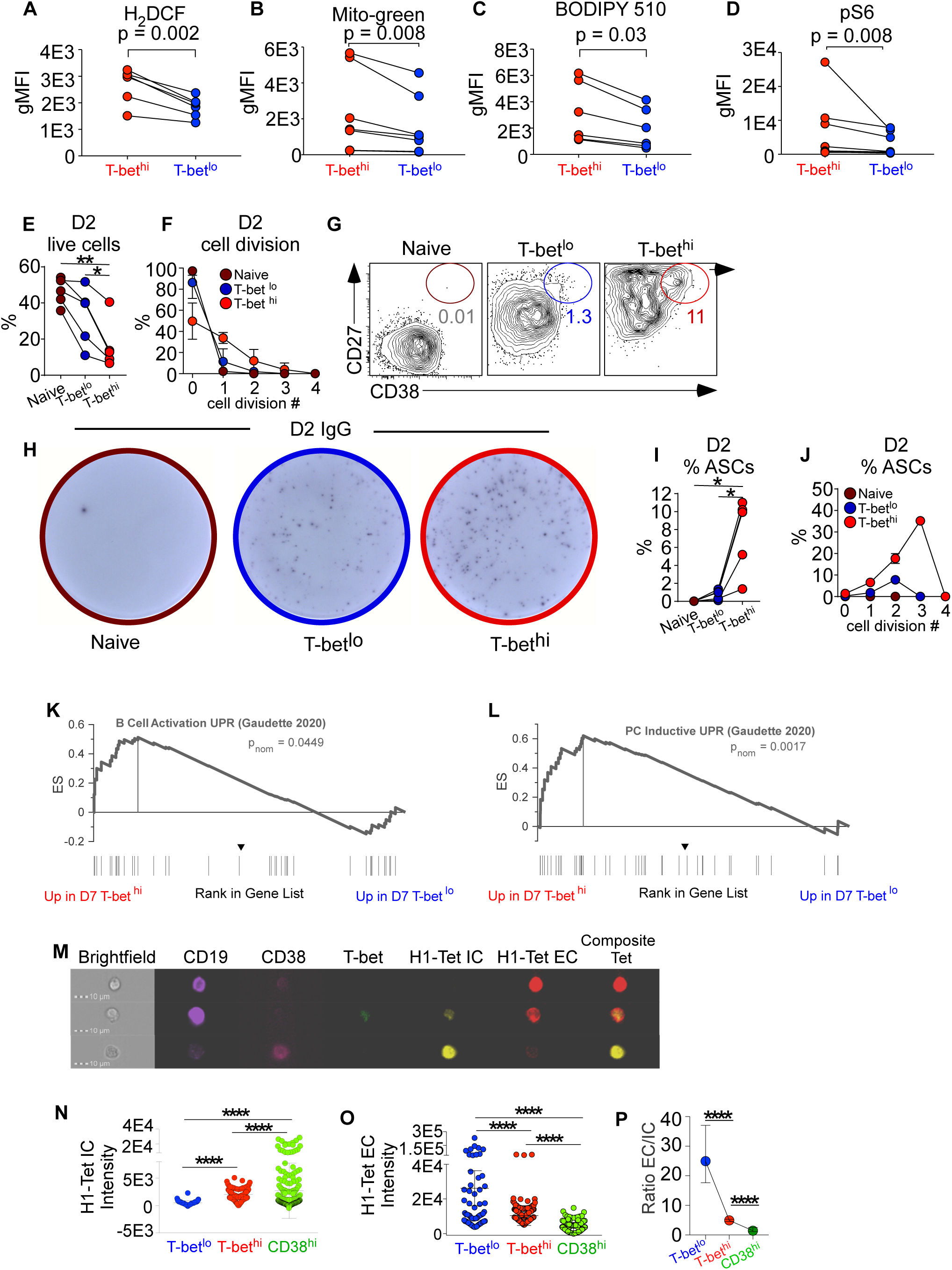
T-bet^hi^ effector memory B cells exhibit metabolic changes associated with ASCs and rapidly differentiate into ASCs. **(A-D)** Metabolic analyses of tonsil-derived donor-matched (n=6-8 donors) T-bet^hi^ (FcRL5^+^, red) and T-bet^lo^ (FcRL5^neg^, blue) IgD^neg^CD27^+^ memory B cells. See Fig. S4A-C for representative gating strategy and Fig. S4D-G for representative flow plots showing metabolic activity in the memory B cell subsets. Data in A-D shown as paired analysis between T-bet^hi^ and T-bet^lo^ cells isolated from the same donor. (A) ROS levels reported as geometric MFI (gMFI) after staining with 2, 7–dichlorohydrofluorescein diacetate (H_2_DCF). (B) Mitochondrial mass reported as gMFI following staining with Mitotracker Green. (C) Membrane lipid analysis reported as gMFI following staining with the fatty acid analog BODIPY510. (D) mTORC1 activation reported as gMFI following staining with anti-phospho-S6 (pS6) kinase Ab. (**E-J**) Donor matched naïve (brown), T-bet^hi^ (FcrL5^+^, red) or T-bet^lo^ (FcrL5^neg^, blue) IgD^neg^CD27^+^CD38^lo/med^ memory B cells were sort-purified (see Fig. S4A-C for sorting strategy), labeled with CTV, stimulated with R848, IL-21, IL-2 and IFN-γ and assessed on D2 (**E-J**) or D3 (Fig. S4H-M). Data shown as the mean ± SD (n= 5 donors, **E, I** and n = 3 donors, **F, J**). **(E-F)** Cell survival (**E**) and frequency of cells in each cell division (**F**) determined by flow cytometry. **(G-J)** ASCs in cultures enumerated by flow cytometry (**G**) and ELISPOT **(H)**. Frequencies of total CD27^hi^CD38^hi^ ASCs (**I**) from each matched culture and percentage of ASCs in each cell division (**J**) are shown. **(K-L)** GSEA comparing the RNA-seq ranked gene list from D7 H1^+^ IgD^neg^ T-bet^hi^ and T-bet^lo^ memory B cells to genes upregulated by the mTORC1-dependent early UPR in mouse follicular B cells (Gaudette et al., 2020) (Table S4) activated with CpG (**K**, B cell activation UPR) or CpG, IL-4 and IL-5 (**L**, PC inductive UPR). Heat map of UPR genes shown in Fig. S4N. **(M-P)** Image Stream analysis of CD38, intracellular T-bet and intracellular (H1-PE tetramer^+^) or extracellular H1 (H1-APC tetramer^+^) expression levels by D7 post-IIV CD19^+^IgD^neg^ cells (see methods). Individual CD38^hi^ PBs (n=211 cells), H1 tetramer-binding T-bet^hi^ (n=195 cells) and T-bet^lo^ (n=62 cells) cells were analyzed. Representative images shown in (**M**). The intensity of intracellular (IC) H1 tetramer staining **(N)** and extracellular (EC) H1 tetramer staining **(O)** and the ratio of EC/IC H1 tetramer staining (**P**) in each T-bet^lo^, T-bet^hi^ and CD38^hi^ cell was determined. Ratio in (**P**) shown as median for the individual cells within each subset and error bars indicate the 95% confidence interval. Statistical analyses were performed using Wilcoxon matched-pairs signed rank tests (**A-D**), one-way ANOVA with Kruskal-Wallis Multiple Comparisons Testing (**N,O**) and one-way ANOVA with Tukey’s multiple comparison testing (**E, I, P**). *p< 0.05, **, p<0.01, *** p<0.001, **** p <0.0001 ns= non-significant

B cell differentiation into ASCs, which can secrete 10,000 Ab molecules per second per cell (van Anken et al., 2003), requires the cells to undergo major changes in metabolism to support the production, secretion and recycling of macromolecules and proteins. Increased protein production, supported by oxidative phosphorylation (Price et al., 2018) and mTORC1 activity (Benhamron et al., 2015) must be ramped up, and compensatory adaptations that allow the cells to cope with increased ER stress, including ROS detoxification (Bertolotti et al., 2012; Pengo et al., 2013) and the unfolded protein response (UPR, (Oikonomou and Hendershot, 2020)) , must be engaged. These adaptations are important as inhibition of these steps is sufficient to prevent development of fully functional ASCs (Shaffer et al., 2004). Given our data showing that the tonsil T-bet^hi^ CD27^+^IgD^neg^ memory B cells exhibit metabolic changes consistent with the necessary ASC adaptations, we predicted that these memory B cells would be poised to differentiate more rapidly into ASCs following re-activation. To test this hypothesis, we stimulated CellTrace Violet (CTV)-labeled, sort-purified matched naïve B cells and IgD^neg^CD27^+^ T-bet^hi^ (FcRL5^+^) and T-bet^lo^ (FcRL5^neg^) memory B cells with TLR7/8 ligand, IFNγ, IL-2 and IL-21 and measured proliferation, survival and ASC development over 72 hrs. As expected, ∼50% of the naïve B cells died in culture during the first two days (Fig. 6E) and the remaining cells (>95%) did not divide (Fig. 6F) or differentiate into ASCs (Fig. 6G-I). Consistent with the increased apoptotic transcriptional signature identified in the T-bet^hi^ vaccine-specific memory B cells, fewer T-bet^hi^ memory cells survived over two days in culture when compared to the T-bet^lo^ memory B cells isolated from the same donor (Fig. 6E). However, in contrast to the naïve and T-bet^lo^ cells, the surviving T-bet^hi^ memory B cells divided ≥1 time within 48 hrs (Fig. 6F). Moreover, while few ASCs were detected in the day 2 cultures containing T-bet^lo^ memory B cells, ASCs were easily measured in the matched day 2 T-bet^hi^ memory B cell cultures (Fig. 6G-I). These ASCs, which were detected by flow cytometry (Fig. 6G, I) and ELISPOT (Fig. 6H), represented more than 50% of the cells that had divided at least once (Fig. 6J). By day 3, some of the naïve B cells had divided (Fig. S4H-I) but ASCs were not detected (Fig. S4J-M). By contrast, >50% of the T-bet^lo^ memory B cells had divided (Fig. S4I) and ASCs were present (Fig. S4J-M) and concentrated within the proliferating fraction (Fig. S4M). Despite this, the T-bet^hi^ memory B cell cultures still contained a significantly higher fraction of ASCs than the matched cultures containing T-bet^lo^ memory B cells (Fig. S4J-M). These data indicate that T-bet^hi^ memory B cells preferentially and rapidly enter cell cycle and differentiate into short-lived ASCs.

### Vaccine-specific T-bet^hi^ memory B cells produce intracellular Ig

Our data showed that bulk T-bet^hi^ CD27^+^IgD^neg^ memory B cells exhibit key metabolic attributes of effector memory cells that appear to poise the cells to rapidly divide, differentiate and secrete Ab. It is appreciated that these early metabolic changes associated with Ab production begin before upregulation of the ASC commitment TFs (Benhamron et al., 2015) and before induction of the XBP1-directed UPR (van Anken et al., 2003). Recent studies (Gaudette et al., 2020) revealed that UPR genes associated with B cell activation (B cell activity UPR) and initiation of the ASC program (PC inductive UPR) were upregulated in mouse follicular B cells activated *in vitro* with TLR ligands and cytokines. These UPR transcriptional programs were induced in an XBP1-independent fashion and were engaged before expression of the ASC lineage TF, BLIMP1. Interestingly, mTORC1 was required for these early XBP1- independent changes in UPR gene expression as well as subsequent differentiation into BLIMP1-expressing ASCs (Gaudette et al., 2020). Given our data showing increased mTORC1 activity in the tonsil T-bet^hi^ memory B cells (Fig. 6D) and the enhanced expression of mTORC1-regulated protein synthesis and degradation pathway genes in the vaccine-elicited HA^+^ T-bet^hi^ memory B cells (Fig. 5D), we predicted that the mTORC1-directed UPR program would be upregulated in the T-bet^hi^ vaccine-induced memory population. To test this hypothesis, we performed GSEA using the B cell activity and PC inductive UPR gene lists ((Gaudette et al., 2020), Table S4) and the ranked gene list from D7 H1^+^IgD^neg^ T-bet^hi^ and T-bet^lo^ memory B cells. Consistent with the effector-like properties of the T-bet^hi^ cells, we observed significant enrichment of both the B cell activity and PC inductive UPR genes in the T-bet^hi^ memory B cells (Fig. 6K-L, Fig. S4N). These data therefore support the conclusion that the D7 vaccine-induced T-bet^hi^ memory B cells have undergone mTORC1-dependent adaptations that support increased protein production associated with the transition to ASCs.

Another feature associated with the transition to an ASC is the switch from producing membrane-associated extracellular (EC) Ig to producing secretory Ig that can be detected in intracellular compartments (IC). This switch is progressive and begins before secretion of Ab and even before expression of BLIMP1 (van Anken et al., 2003). Using ImageStream and H1 tetramers in two different fluorochromes, we quantified the IC and EC forms of Ig in the memory subsets and CD38^hi^ PBs (Fig. 6M). As expected, PBs expressed abundant and easily detected IC H1-specific Ig (Fig. 6N), which was accompanied by a large reduction in EC H1-specific Ig (Fig. 6O). The T-bet^lo^ memory B cells expressed high levels of EC H1-specific Ig and low to undetectable levels of IC H1-specific Ig (Fig. 6N-O). In contrast, T-bet^hi^ H1-specific memory T cells reproducibly expressed lower levels of EC H1-binding Ig than the T-bet^lo^ H1^+^ subset (Fig. 6O). Strikingly, T-bet^hi^ effector memory cells also expressed easily measurable levels of IC H1-specific Ig (Fig. 6N). Moreover, the EC/IC H1-binding Ig ratio in individual cells from each population (Fig. 6P), revealed that the T-bet^hi^ vaccine-specific memory B cells were more similar to the PBs than to the T-bet^lo^ memory B cells. Taken together, these data show that the IgD^neg^ HA-specific T-bet^hi^ memory B cells found in circulation after vaccination exhibit the metabolic and functional attributes of a poised Ig producing effector memory B cell.

### T-bet^hi^ memory B subset contributes to secondary ASC responses and predicts enduring humoral immunity to IIV

T cell effector memory populations, particularly those localized in tissues, appear to decline more rapidly than memory cells with stem-like properties (Woodland and Kohlmeier, 2009). Our data showed that clonotypes from the T-bet^hi^ effector memory population were detectable in blood for at least one-month post-vaccination (Fig. 3E). To assess whether vaccine-elicited T-bet^hi^ effector population is short or long-lived, we immunized 11 HD, who self-declared as missing prior annual flu vaccines, over two sequential vaccine seasons with flu vaccines that contained the same H1 or H3 Ag both years (Fig. 7A). On D7 post-vaccination each year, we determined the frequency of T-bet^hi^ and T-bet^lo^ memory B cells specific for the repeated HA Ag. On D7 post-immunization with the 2015 influenza vaccine (Fig. 7A), we identified both T-bet^hi^ and T-bet^lo^ CA09-H1 specific B cells (Fig. 7B). Following revaccination of those individuals in 2016 with the same CA09-H1 Ag, we observed a decline in the CA09-H1 reactive T-bet^hi^ B cell response, relative to the response made by that individual in 2015 (Fig. 7B). However, the frequencies of CA09-H1-specific T-bet^lo^ memory B cells were similar in both years (Fig. 7B). Similarly, when HD were immunized in 2016 with the HK14-H3-containing vaccine (Fig. 7A), we identified HK14-H3-specific T-bet^hi^ and T-bet^lo^ memory B cells (Fig. 7C). Following revaccination in 2017 with the same HK14-H3 Ag, we observed a decrease in the fraction of T-bet^hi^ HK14-H3^+^ memory B cells with no concomitant loss of the T-bet^lo^ population (Fig. 7C). When we compiled the reactivity data to sequentially administered HA Ags across all 11 donors, we observed no difference in the frequencies of circulating T-bet^lo^ vaccine-specific memory B cells over 2 years (Fig. 7D) but a significant reduction in the T-bet^hi^ vaccine-specific memory B cell subset between year 1 to year 2 (Fig. 7D).

**Figure 7.**
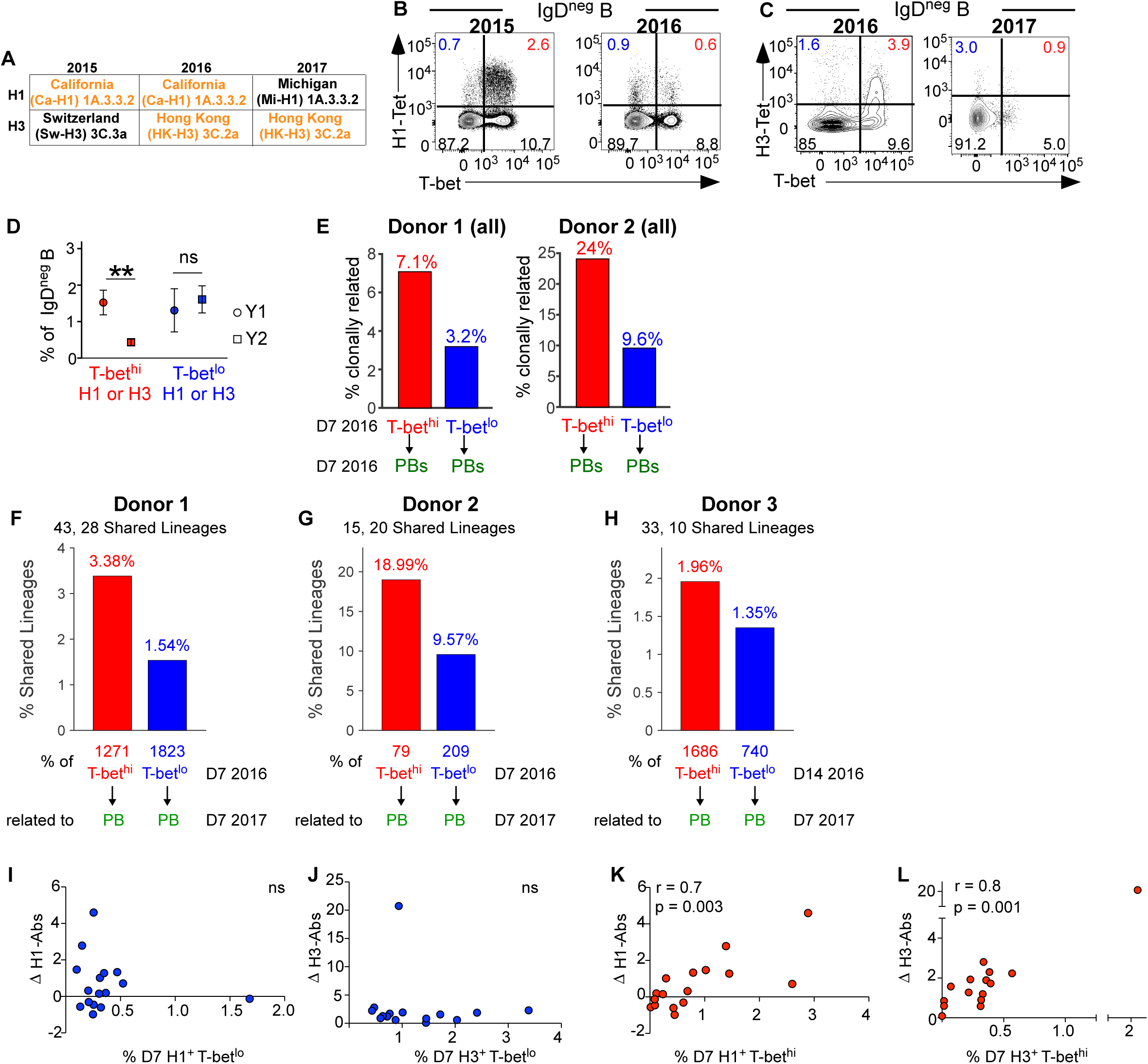
HA-specific IgD^neg^ T-bet^hi^ effector memory B cells contribute to secondary ASC responses and predict enduring humoral immunity. **(A)** Composition of H1 and H3 Ags in 2015-2017 IIV. (**B-D**) Enumeration of D7 H1- or H3-specific IgD^neg^ T-bet^hi^ and T-bet^lo^ B cells in 11 HD sequentially immunized with IIV between 2015 and 2016 (n=6) or between 2016 and 2017 (n=5). **(B)** Representative FACS plots showing H1-CA09^+^ memory B cells over 2 years (2015-2016) from a single individual immunized with IIV containing the same CA09-H1 Ag each year. **(C)** Representative FACS plots showing HK14-H3 memory B cells over 2 years (2016-2017) from a single individual vaccinated with IIV containing the same HK14-H3 Ag each year. Frequencies of T-bet^hi^ and T-bet^lo^ B cells within the H1- or H3-specific IgD^neg^ population indicated. **(D)** Frequency, represented as mean with standard error, of the H1^+^ or H3^+^ T-bet^hi^ (red) or T-bet^lo^ (blue) B cells for 11 HD vaccinated over 2 years with IIV containing matched H1 or H3 Ags. ** p = 0.002 Wilcoxon matched-pairs signed rank tests. **(E)** IgH(V_H_) BCR repertoire analysis (Table S1E-F) to identify shared lineages between PBs and H3^+^ IgD^neg^ T-bet^hi^ and T-bet^lo^ B cells isolated from HD (n=2) on D7 post-IIV (2016). Data reported as percentage of shared lineages with the number of shared lineages between PBs and T-bet^hi^ cells and between PB and T-bet^lo^ cells (top) and number of non-singleton lineages identified in each subset (bottom) shown. Connections between lineages in Fig. S5A. (**F-H**) IgH(V_H_) BCR repertoire analysis (Table S1E-F) performed on sorted D7 H3^+^ IgD^neg^ T -bet^hi^ and T-bet^lo^ memory B cells (**F**-**G**) or D14 H1^+^ IgD^neg^ T -bet^hi^ and T-bet^lo^ memory B cells (**H**) and compared to IgH(V_H_) BCR repertoire analysis from sorted D7 PB following re-vaccination of the same donor the following year (2017). Data are shown as percentage of shared lineages between the 2016 T-bet^hi^ or T-bet^lo^ subsets and the 2017 PB with the number of shared lineages (top) and number of non-singleton lineages identified in each subset (bottom) indicated. (**I-L**) Assessment of vaccine-specific D7 memory B cell responses (frequency in blood) and vaccine-specific Ab responses (measured as FC in titers between D0 and D120) after IIV (2015) in HD (n=19). (**I-J**) Correlation between FC in H1- (**I**) and H3- (**J**) specific IgG titers (Y axis) and the frequency of the D7 H1- (**I**) and H3- (**J**) specific T-bet^lo^ cells within the IgD^neg^ B subset (X axis). (**K-L**) Correlation between the FC in H1- (**K**) and H3- (**L**) specific IgG titers (Y axis) and the frequency of the D7 H1- (**K**) and H3- (**L**) specific T-bet^hi^ cells within the IgD^neg^ B subset (X axis). ns = no significant correlation. Spearman correlation (r) and significance (p) between D7 memory B cell subsets and FC in Ab responses are indicated. Additional correlation analyses for D7 and D14 T-bet^hi^ and T-bet^lo^ vaccine-specific memory B cells shown in Fig. S5B-H.

Our data showed that the circulating T-bet^hi^ memory B cell subset was greatly reduced after re-exposure to the same Ag one year later. These data were consistent with the possibility that the effector memory population is relatively short-lived. Alternatively, the long-lived T-bet^hi^ effector memory population might be diverted from the memory compartment into the early PB response following Ag re-exposure. To test this possibility, we first performed BCR repertoire analysis to determine whether D7 PBs were more clonally related to the D7 T-bet^hi^ or T-bet^lo^ IgD^neg^ H3^+^ B cells circulating in the same vaccinated individuals (Table S1E-F). We observed that 7% of the D7 T-bet^hi^ lineages were also found in the circulating D7 PBs from the same donor (Fig. 7E). The shared lineages between these two populations included both large and small clonotypes (Fig. S5A). By contrast, only 3% of the T-bet^lo^ lineages were shared with the D7 PBs from the same donor (Fig. 7E). Similar results were seen with a second donor (Fig. 7E, Fig. S5A). These data were consistent with the idea that T-bet^hi^ HA^+^ effector memory cells that are recalled to highly related Ags might enter the ASC compartment. To further assess this possibility, we sorted the T-bet^hi^ and T-bet^lo^ HA-specific memory B cells on D7 or D14 from three different HD and then reimmunized those HD a year later with a vaccine containing the same H3 Ag and a very similar H1 Ag (97.4% identity). We isolated the circulating PBs on D7 following the repeat vaccination in year 2 and then compared the BCR repertoire between the year 1 memory B cells and year 2 PBs. Consistent with our hypothesis, more lineages were shared between the year 1 T-bet^hi^ memory B cells and the year 2 PBs relative to what we observed when we compared lineage sharing between the Year 1 T-bet^lo^ memory B cells and the year 2 PBs from the same individuals (Fig. 7F-H). These data therefore suggest that at least some of the clonotypes within the vaccine-induced, circulating T-bet^hi^ memory B cell subset persist and contribute to the rapid memory B cell-derived ASC recall response.

Our data suggested that at least some of the cells within the T-bet^hi^ memory B cell subset are long-lived and might therefore be derived from GC reaction (Takahashi and Kelsoe, 2017). If so, the T-bet^hi^ memory B cell population should correlate with other GC-dependent events, including the development of the long-lived systemic Ab response, which can be discriminated from the transient IIV-induced Ab response that declines within ∼90 days (Nakaya et al., 2015). To test this hypothesis, we determined the FC in the vaccine-specific H1 and H3 IgG Ab response between day 0 and day 120 post-immunization in 19 vaccinated HD and compared this response to the size of the D7 PB, T-bet^hi^ and T-bet^lo^ H1-specific or H3-specific memory B cell responses in those individuals. As previously reported (Wrammert et al., 2008), the early circulating PB response did not correlate with the long-lived humoral immune response generated in that individual (Fig. S5B). Similarly, we observed no correlation between the size of the T-bet^lo^ H1 (Fig. 7I) or H3 (Fig. 7J) responses on either D7 (Fig. 7I-J) or D14 (Fig. S5C-D) post-vaccination and the change in systemic Ab titers 4 months after vaccination. However, we observed a strong positive correlation between the size of the T-bet^hi^ H1 (Fig. 7K) and H3 (Fig. 7L) specific memory B cell subset on D7 (Fig. 7K-L) and D14 (Fig. S5E-F) and the magnitude of the long-lived H1 or H3 specific humoral immune response generated by vaccination. Importantly, this was Ag dependent as no correlation was seen between the T-bet^hi^ *<u>H1</u>*-specific memory B cell response and the change in *<u>H3</u>* Ab titers or vice versa (Fig. S5G-H). Taken together, the molecular, cellular and functional data each support the conclusion that the T-bet^hi^ CD27^+^ memory B cell subset, which can be found in human tissues and in circulation after vaccination, represent a population of poised Ig producing “effector” human memory B cells that can contribute rapidly to secondary ASC responses following reactivation. In addition, these cells may represent a rapid, easily-measured circulating biomarker of GC responses that can be used to predict whether an individual is likely to generate a long-lived humoral response to seasonal IIV.

## Discussion

Immunization with seasonal IIV predominantly recruits and recalls memory B cells that were generated in response to prior infections or immunizations (Andrews et al., 2015; Turner et al., 2020). The IIV-elicited memory B cell compartment is composed of multiple subsets that reflect their different origins, molecular properties and functional attributes. Prior studies analyzing IIV-specific IgD^neg^ Ag-experienced B cells identified at least two populations of recirculating memory B cells, which were defined as CD27^+^CD71^neg^ resting memory (RM) cells and CD71^+^ activated memory (AM) B cells (Ellebedy et al., 2016). The vaccine-specific AM compartment was subsequently further subdivided into the IgD^neg^CD27^neg^ population (Andrews et al., 2019) and the IgD^neg^CD27^+^ subset, which includes both CD21^lo^ and CD21^hi^ cells (Andrews et al., 2019; Koutsakos et al., 2018; Lau et al., 2017). The CD27^+^CD21^lo^ memory compartment is reportedly enriched in transcriptionally programmed BLIMP-1 expressing pre-ASCs (Lau et al., 2017) while the IgD^neg^CD27^neg^ subset represents newly-formed, primary memory B cells that arise from the GC reaction (Andrews et al., 2019). Our data adds to this compendium as we characterize a population of vaccine-induced IgD^neg^CD27^+^ T-bet expressing human “effector-like” memory B cells that, while not expressing ASC fate-determining TFs, are metabolically poised to rapidly assume the functional properties of ASCs after reactivation.

Based on our previous animal model data showing that T-bet is required for memory B cell-derived secondary ASC responses (Stone et al., 2019), we predicted that T-bet might serve as a useful biomarker to subdivide human memory B cells into distinct subsets that would differ in their transcriptional programming and presumably their functional attributes. Using extracellular FcRL5 staining as a faithful surrogate for intracellular T-bet expression to subdivide IIV-elicited HA-specific memory cells, we characterized two discrete IgD^neg^CD27^+^ memory populations, which included the T-bet^lo^FcRL5^neg^ memory cells and T-bet^hi^FcRL5^+^ memory cells. Based on isotype usage, accumulation of somatic mutations, and the presence of high avidity and broadly reactive cells, it is clear that both subsets include Ag-experienced *bona fide* memory cells. Moreover, clonotypes from both subsets can be recalled into secondary responses, indicating that each subset contributes to the long-lived memory compartment. The T-bet^lo^ cells, which are CD71^neg^CXCR5^+^ CCR7^+^CD62L^+^, are phenotypically similar to the previously described RM B cell compartment (Andrews et al., 2019; Ellebedy et al., 2016) and transcriptional profiling of these cells revealed that they express higher levels of *TCF7* and *BACH2 -* TFs that are associated with increased proliferation potential and stem-like qualities (Pais Ferreira et al., 2020; Tsukumo et al., 2013; Yao et al., 2021; Zhou et al., 2010). Thus, the T-bet^lo^ memory population appears to be enriched for resting stem-like memory cells.

Although the D7 T-bet^hi^ HA-specific memory B cells share some phenotypic and transcriptional attributes with other previously described vaccine-elicited B cell populations, including the CD27^neg^IgD^neg^ HA-specific primary memory B cells that arise late after vaccination (Andrews et al., 2019) and the CD27^+^CD21^lo^ pre-ASCs that are found 14 days after vaccination (Lau et al., 2017), we believe that the D7 T-bet^hi^FcRL5^+^ CD27^+^ memory population represents a molecularly distinct population that is most similar to the AM2 population that was identified, but not analyzed in depth, in individuals receiving a prime/boost H7N9 vaccine (Andrews et al., 2019). Transcriptional and epigenetic profiling of the CD27^+^CCR7^neg^CXCR5^neg^CD62L^neg^ HA-specific T-bet^hi^ memory B cells reveals that these cells share considerable overlap with effector and effector memory CD8 T cells. The T-bet^hi^ effector memory subset also expresses increased levels of transcriptional regulators like *TOX2*, *EZH2*, and *ARID3A* that are constitutively expressed by ASCs. Given that ASCs represent the terminally differentiated B lineage effector cell type we concluded that the HA-specific T-bet^hi^ CD27^+^ cells might represent an effector-like memory population.

Effector CD8 T cells can be distinguished from central memory T cells based on changes in cellular metabolism, respiration and mitochondrial dynamics (Buck et al., 2017). Consistent with this, the T-bet^hi^ effector-like memory B cells express increased levels of genes linked to numerous metabolic processes that support energy intensive processes like protein and macromolecule synthesis. In order to combat increased ER stress that arises as a result of the increased metabolic activity, the T-bet^hi^ cells have also upregulated anti-oxidant defense systems and genes associated with ER-proteasome and ER-autophagosome pathways of protein degradation and recycling (Oikonomou and Hendershot, 2020). Importantly, many of the key changes in transcriptional networks associated with mitochondrial activity, oxidative stress, fatty acid synthesis and mTORC1 activity are recapitulated in functional studies using the phenotypically similar T-bet^hi^ FcRL5^+^ CD27^+^ tissue-residing memory B cells. Similar metabolic programming changes are seen in ASCs (Lam et al., 2018) and are, in fact, required for the development of fully functional ASCs (Lam and Bhattacharya, 2018). Thus, the T-bet^hi^ memory B cells exhibit metabolic characteristics that are associated with ASCs.

It is not surprising that terminally differentiated effector ASCs, which secrete Ab at truly astonishing rates, must undergo extensive metabolic and structural changes, such as switching from glycolytic metabolism to oxidative phosphorylation (Price et al., 2018) and utilizing other energy sources, like fatty acids and glutamine (Lam and Bhattacharya, 2018). One critical aspect of these metabolic changes is that the cells must adjust to the ER stress that arises as a consequence of misfolded proteins that are inevitably generated as protein synthesis rates increase. This process (Oikonomou and Hendershot, 2020), referred to as the unfolded protein response (UPR), activates cellular redox reactions and targets misfolded proteins to the proteasome or to the lysosome/autophagosome complex through Endoplasmic Reticulum Associated Degradation (ERAD) and ER-phagy reactions (Oikonomou and Hendershot, 2020). In ASCs, the UPR is induced and maintained by the TFs IRE1, XBP1 and ATF6 (Oikonomou and Hendershot, 2020) and is associated with the downregulation of PERK, the PKR-like ER kinase (Ma et al., 2010). Despite the fact that the T-bet^hi^ effector memory B cells have downregulated the PERK gene (*EIF2AK3*) and express higher levels of multiple UPR and ERAD-associated genes, like *OS9*, *UBA1* and *VCP*, these cells do not express increased levels of *XBP1*, *IRE1* or *ATF6*. These data therefore suggest that the T-bet^hi^ memory B cells regulate cellular stress responses through a XBP1- independent mechanism.

Metabolic changes, particularly those that support increased secretory Ig synthesis, are initiated before ASC committing TFs like BLIMP1 are induced (Benhamron et al., 2015) and well before expression of XBP1 (van Anken et al., 2003), which is not obligate for ASC development (Iwakoshi et al., 2003; Taubenheim et al., 2012). A recent publication (Gaudette et al., 2020) reported that UPR-associated genes are upregulated following B cell activation and before ASC commitment and showed that this process is XBP1-independent. Instead, the induction of the UPR in these cells requires mTORC1 (Gaudette et al., 2020) – a key metabolic sensor and driver of protein synthesis (Saxton and Sabatini, 2017) and a known regulator of B cell differentiation and function (Raybuck et al., 2019). The mTORC1-dependent UPR gene signature is significantly enriched in the transcriptome of the T-bet^hi^ memory B cells, indicating that this protective mechanism is likely constitutively active in these cells. This is consistent with the data showing increased expression of mTORC1 complex genes, like LAMTOR1, LAMTOR2 and LAMTOR5, in the T-bet^hi^ memory B cell subset and increased basal mTORC1 activity in these cells. Thus, the modified metabolism of the T-bet^hi^ memory subset could both facilitate increased protein synthesis and engage the compensatory programs necessary to support increased protein production.

Like the effector memory T-bet^hi^ B cell subset, UPR genes are basally expressed in the marginal zone B cell subset (Gaudette et al., 2020). Marginal zone B cells represent a highly responsive population that rapidly differentiates into ASCs following activation (Gunn and Brewer, 2006). Similarly, *in vitro* stimulated T-bet^hi^ memory B cells enter cell cycle and differentiate into antibody producing ASCs more quickly than the T-bet^lo^ memory cells. The accelerated secretory response of the T-bet^hi^ memory B cells is likely due to their ongoing production of intracellular Ig. Indeed, the ratio of plasma membrane-associated Ig to intracellular Ig is already quite similar between the HA-specific T-bet^hi^ memory subset and ASCs. Interestingly, examination of secondary ASC responses in three re-vaccinated individuals indicate that the clonotypes from the T-bet^hi^ memory subset are more highly represented in the early D7 secondary ASC response than clonotypes from the T-bet^lo^ subset. This result does not mean that the T-bet^lo^ memory subset cannot contribute to secondary ASCs responses. In fact, T-bet^lo^ memory B cell clonotypes are present, albeit at lower percentages, in secondary ASCs. Rather, the data are consistent with the T-bet^hi^ memory B compartment being metabolically poised to rapidly initiate their effector activities upon reactivation and therefore able to contribute more rapidly to the ASC pool.

Our data showing that T-bet expression within the murine memory B cell compartment is important for secondary ASC responses to viral infection (Stone et al., 2019) suggests that T-bet plays an active role in effector memory development or function. Although we cannot definitively answer whether T-bet is also important for the vaccine-elicited T-bet^hi^ effector-like human memory B cells, the data support a role for T-bet in regulating chromatin accessibility within the genome of the T-bet^hi^ memory population. Consistent with this, T-bet is associated with epigenetic changes in specific pathways and genes that could influence the properties of the T-bet^hi^ memory cells. For example, IPA predicts BCR signaling as the top differentially regulated pathway between T-bet^hi^ and T-bet^lo^ memory B cells when the analysis is restricted to genes linked to DAR containing T-bet binding motifs. This suggests that, through its capacity to control chromatin accessibility, T-bet might poise BCR signaling genes to rapidly respond to external cues. Alternatively, T-bet may function as a master regulator by influencing expression of other transcriptional regulators that are important for preserving the memory program or preparing the cells to initiate new effector functions. In fact, *BACH2*, which must be downregulated during the commitment to the ASC lineage (Igarashi et al., 2014; Kometani et al., 2013), exhibits a loss of chromatin accessibility surrounding a previously identified T-bet binding site. By contrast, chromatin accessibility surrounding known T-bet binding sites is increased within the *TOX2* and *ARID3A* loci of the T-bet^hi^ memory B cells. Given that TOX2 supports chromatin accessibility (Xu et al., 2019) and ARID3A is a transactivator of IgH locus (Lin et al., 2007) these data suggest that T-bet likely contributes to the epigenetic changes that support the fate decisions and effector functions of the T-bethi effector memory B cell subset.

We observed that the size of the D7 post-vaccination T-bet^hi^ memory B cell response correlated significantly with the development of an enduring long-lived serum Ab response (>120 days). This is important as *easily-measured rapid* surrogates that predict long-lived humoral immunity to vaccination are lacking (Nakaya et al., 2015). Indeed, the early circulating PB and T_FH_ cells which, while good surrogates for predicting vaccine responsiveness, do not predict the durability of the vaccine-induced humoral response (Bentebibel et al., 2013; Nakaya et al., 2015). Our data suggest that the T-bet^hi^ memory response may be linked to the same types of GC-centric processes that generate the long-lived PC compartment. While data in mice suggest that memory B cells may not contribute significantly to GC responses (Mesin et al., 2020), emerging evidence in humans indicates that both naïve and memory B cells participate in LN GC responses following IIV (Turner et al., 2020). Our data, which show that the T-bet^hi^ memory B cell clonotypes are high avidity, extensively mutated and can be recalled into secondary vaccine-elicited ASC responses at least one year after the first vaccination, support the conclusion that these cells have, at some point in their ontogeny, undergone GC selection. Interestingly, despite the fact that stem-like T-bet^lo^ HA^+^ memory cells are also long-lived and appear to have undergone GC selection, the size of this population does not correlate with the long-lived PC response. This suggests that the appearance of the HA-specific T-bet^hi^ and T-bet^lo^ populations in circulation following vaccination may be controlled by different signals. While we cannot say with certainty that the responding T-bet^hi^ memory cells are recent GC emigrants, we can say that the rapid emergence of this HA-specific memory population accurately predicts new GC-dependent long-lived PC responses. Thus, the IIV-elicited T-bet^hi^ effector-like memory subset is not only important as a pool of memory cells that can quickly contribute a new burst of secreted Ab following re-exposure to the same or closely related Ags but may be practically useful as easily monitored rapidly induced population that can be used to predict the development of a long-lived Ab response to vaccination.

## Author Contributions

F.E.L. conceived the idea for the project and secured the initial funding. F.E.L. and A.N. designed the experiments that were performed by A.N., E.Z., C.D.S., R.G.K, C.M.T., B.M. and F.Z. B cell tetramers were developed and produced by J.E.B. Human samples used in this study were obtained via the Alabama Vaccine Research Clinic, directed by P.A.G. Bioinformatic analyses were performed by A.F.R, C.D.S., C.F. and T.M. All other data was analyzed by A.N and F.E.L. A.N., A.F.R. and F.E.L wrote the manuscript and prepared final figures. Critical feedback on the project and manuscript were provided by A.F.R, T.D.R, J.F.K., I. S., and J.M.B.

## Acknowledgements

We thank the Alabama Vaccine Research Clinic and Pamela Cunningham, Heather Logan, Aeryn Peck, Catrena Johnson and Megan Oelschig for recruiting and consenting subjects for this study. We thank Vidyasagar Hanumanthu, Director of the UAB Flow Core Facility, for his assistance with flow sorts and ImageStream experiments and Dr. Michael Crowley, Director of the UAB Heflin Genomics core, for his assistance with next-gen sequencing. We are grateful to Dr. Amy S. Weinmann for her thoughtful commentary regarding this manuscript. Funding for the work was provided by the US National Institutes of Health (NIH): P01 AI125180 (I.S., F.E.L, J.M.B, C.D.S, E.Z.), U19 AI 109962 (F.E.L, T.D.R) and U19AI142737 (F.E.L., T.D.R., A.N., R.D.K., A.R.) A.N. received salary support from R21 AI152006, pilot grant support from the UAB AMC21 Immunology Autoimmunity and Transplantation Strategic Initiative as well as funding from the UAB CCTS (UL1 TR001417) to support this work. C.D.S. received salary support from R01 AI148471. NIH P30 AR048311 and P30 AI027767 provided support for the UAB Consolidated Flow Cytometry Core and NIH P30CA13148 provided support for the O’Neal Comprehensive Cancer Center Tissue Procurement Facility. The authors have no known conflicts of interest to disclose.

## Experimental Procedures

### Human Subjects and Samples

The UAB Institutional Review Board approved all human study protocols. Subjects, who self-identified as healthy, were recruited and consented through the Alabama Vaccine Research Clinic (AVRC). Subjects received 2015-2016 Fluzone (Sanofi-Pasteur), 2016-2017 Fluvirin (Sequiris), 2017-2018 Fluzone (Sanofi-Pasteur) or 2018-2019 Fluzone (Sanofi-Pasteur). Blood was drawn on days 0, 7, 14, 21, 28 and 120 days +/- 1 week. The O’Neal Comprehensive Cancer Center Tissue Procurement Shared Facility provided remnant tonsil tissue samples from patients undergoing routine tonsillectomies.

### Lymphocyte and plasma isolation

Peripheral blood from human subjects was drawn into K2-EDTA tubes (BD Bioscience). Peripheral blood mononuclear cells (PBMCs) and plasma were isolated by density gradient centrifugation over Lymphocyte Separation Medium (CellGro). Red blood cells were lysed with ammonium chloride solution (StemCell). Plasma and PBMCs were either used immediately or aliquoted and stored at -80°C.

### Recombinant influenza HA protein production

The coding sequences (amino acids 18-524) of influenza HA ectodomains were synthesized (GeneArt, Regensburg, Germany) from the following influenza virus strains: A/California/VRDL7/2009 (CA09-H1), A/Switzerland/9715293/2013 (SW13-H3), A/Hong Kong/4801/2014 (HK14-H3), and A/Michigan/45/2015 (MI15-H1). As previously described (Allie et al., 2019), HA ectodomains were cloned into the pCXpoly(+) vector that was modified with a 5’ human CD5 signal sequence and a 3’ GCN4 isoleucine zipper trimerization domain (GeneArt) that was followed by either a 6XHIS tag (HA-6XHIS construct) or an AviTag (HA-AviTag construct). The HA–6XHIS and HA–AviTag constructs for each HA were co– transfected using 293Fectin Transfection Reagent into FreeStyle™ 293–F Cells (ThermoFisher Scientific) at a 2:1 ratio, respectively. Transfected cells were cultured in FreeStyle 293 Expression Medium for 3 days and the supernatant was recovered by centrifugation. Recombinant HA molecules were purified by FPLC using a HisTrap HP Column (GE Healthcare) and eluted with 250 mM imidazole.

### Generation of HA tetramer and HA cytometric bead arrays

Recombinant HA trimers were biotinylated by addition of biotin–protein ligase (Avidity). To generate HA tetramers, biotinylated HA trimers were tetramerized with fluorochrome–labeled streptavidin (Prozyme). Labeled tetramers were purified by size exclusion on a HiPrep 16/60 Sephacryl S–300 column (GE Healthcare). To generate cytometric bead arrays (CBA), carboxy functionalized blue particle array beads (Spherotech) were directly conjugated with streptavidin (Southern Biotech) following manufacturer’s protocol. Biotinylated HA tetramers were passively absorbed onto CBA particles by mixing a 5-fold excess of particle binding capacity (100 μg) of biotinylated HA with 2e7 Spherotech beads in 400 μl of PBS with 1%BSA. Labeled microparticles were separated by centrifugation at 3000g for 5min. HA-trimer conjugated beads were washed twice in 1ml of 1% BSA, PBS, 0.05% NaN3, resuspended at 1 x 10^8^ beads/mL and stored at 4°C. IgG capture beads were generated by the direct conjugation of goat anti-Human IgG F(ab)^2^ (Southern Biotech) to array particles (Spherotech) following manufacturer’s protocol for direct labeling. Briefly, 2e7 particles were pelleted by centrifugation and resuspended in 200 μl of anti-human IgG. An equal volume of EDC (1-Ethyl-3-(3-dimethylaminopropyl)-carbodiimide), 6mg/mL, in 0.1 M MES (2-(N-morpholino) ethanesulfonic acid) buffer pH 5.0 was added and the mixture was rotated overnight at RT. Beads were washed twice by pelleting by centrifugation and resuspension in PBS. Following washing, beads were resuspended in 1% BSA in PBS with 0.005% NaN3 as a preservative.

### Flow Cytometry

Single cell suspensions were blocked with 2% human serum and stained with Ab panels described in Table S12. 7AAD or LIVE/DEAD Fixable Dead Cell Stain Kits (Molecular Probes/ThermoFisher) were used to discriminate live cells. To detect HA-binding B cells, cells were treated at 37°C with 0.5U/ml neuraminidase (*C. perfringens*, Sigma) to remove sialic acid, and then were washed, blocked and stained with HA tetramers. Intracellular proteins were detected by staining with Abs specific for cell surface markers, fixing the cells in 10% neutral buffered formalin solution (Sigma), and then staining the permeabilized cells (0.1% IGEPAL (Sigma)) with Abs or fluorochrome-labeled HA tetramers. Stained cells were analyzed using a FACSCanto II (BD Bioscience) or the Attune NxT flow cytometer (Invitrogen, ThermoFisher). FlowJo v9.9.3 or FlowJo v10.2 were used for analysis.

### Cell Sorting

B cell subsets were sort-purified for BCR-seq, RNA-seq, ATAC-seq, rMAb generation, Mitotracker green assays, mTORC1 pS6 assays and *in vitro* culture experiments with a FACSAria (BD Biosciences) or Melody (BD Biosciences) in the UAB Comprehensive Flow Cytometry Core. Table S12 shows flow panels used for subset sort-purifications.

### BCR Cloning and screening of recombinant Abs

B cell subsets were index-sorted (Table S12) and deposited as single cells into hypotonic lysis buffer in 384 well plates that were stored at -80°C. cDNA was generated using High-capacity cDNA generation kit (Roche). PCR amplicons, using forward and reverse primers specific for the amino terminus of the mature IGHV, IGKV, and IGLV proteins (Table S13), were cloned into pEGFP expression vectors that were modified to express the constant regions of human IgG1, IgKappa, or IgLambda. Plasmids encoding Ig heavy and light chain pairs were co-transfected using Polyethylenimine into 293FreeStyle cells (Invitrogen). Supernatants were assayed for secreted rAb and for HA reactivity using the HA CBA described above. The MFI of rAb binding to each HA Ag (CA09, MI15, PR8, HK14) in the bead array was measured by flow cytometry (BD LSRII) and normalized for Ig by dividing the HA binding MFI by the MFI of binding to the anti-IgG capture beads (see Table S1A).

### BCR Repertoire library preparation and informatics

B cell subsets were sorted directly into RLT buffer and snap frozen in LN_2_. RNA was extracted using the quick start protocol from QIAGEN RNeasy Mini Kit. First strain cDNA synthesis was performed using iScript cDNA synthesis kit (BioRad) and 8 μl of RNA. First round amplification of IgG, IgA, and IgM was performed in 25 μl reaction volume using 4-8 μl cDNA, Platinum PCR SuperMix High Fidelity (Invitrogen), and 1μl gene specific primers (120 nM) of VH1-VH7 FR1 (forward) and Ca, Cu, Cg (reverse) (Tipton et al., 2015). First round PCR conditions were: 95°C for 3 min, 42 cycles of 30s 95°C, 30s 58°C, 30s 72°C, and 72°C for 3 min. PCR products were verified on 1.2% agarose gels and then indexed in a 2^nd^ round PCR reaction using the Nextera Index kit (Illumina). PCR2 conditions for indexing were: 72°C for 3 min, 98°C for 30s and 5 cycles of 98°C for 10s, 63°C for 30s, and 72°C for 3 min. Indexed PCR products were purified with Agencourt AMPure XP beads (Beckman), quantitated by NanoDrop and then pooled into libraries. Libraries were denatured using 0.2N NaOH and quenched with cold HT1 per manufacturer (Illumina) instruction. Denatured libraries were diluted with 20% PhiX (Illumina) as an internal quality control and loaded onto a 600-cycle V3 MiSEQ cartridge (Illumina) for amplification.

Ig heavy-chain sequencing data were processed as described previously (Tipton et al., 2015). Briefly, joined paired-end reads were assembled using Fastq-join (https://github.com/ExpressionAnalysis/ea-utils) and quality filtered. Sequences with ≤ 200bp or with low-quality bases (> 0bp with quality scores < 10, or ≥ 5bp with quality scores < 20, or ≥ 15bp with quality scores of 30) were eliminated from further analysis. Overall quality of sequences was assessed using FastQC (https://www.bioinformatics.babraham.ac.uk/projects/fastqc/). Sequences were annotated for isotype and submitted to IMGT/High V-QUEST (Alamyar et al., 2012) web portal for V_H_-, J_H_-gene annotations, alignments and tabulation of mutations. Sequences annotated as “productive” were considered for further analysis. For each donor, sequences (from all B cell subsets) were clustered into lineages based on shared rearrangements (same V_H_ and J_H_ genes, and same HCDR3 length) as well as pairwise HCDR3 nt sequence similarity ≥ 85% (see Supplementary Note 1 in Tipton et al (Tipton et al., 2015)). Mutation frequencies were determined based on non-gap mismatches of expressed sequences relative to their annotated V_H_ germline sequences. Downstream analysis and visualization were performed in Matlab (R2020a, The Mathworks Inc., Natick MA). All BCR sequencing data is available from the Sequence Read Archive (SRA) database under the accession XXXXX. See Table S1.

### RNA-seq library preparation and analysis

B cell subsets were sorted directly into RLT buffer (Qiagen) with 1% mercaptoethanol and then snap-frozen in LN_2_. RNA was extracted using the QuickRNA Micro Prep Kit (Zymo) and cDNA was prepared using SMART-seq v4 cDNA synthesis kit (Takara). Sequencing libraries were constructed using 200 pg cDNA as input for the NexteraXT kit with NexteraXT indexing primers (Illumina). Libraries were quality assessed, pooled and sequenced using 75 bp paired-end chemistry on a NextSeq500 at the UAB Helfin Genomics Core. Sequencing reads were mapped to the hg38 version of the human genome using STAR (Dobin et al., 2013) with the default settings and the UCSC KnownGene table as a reference transcriptome. Reads overlapping exons were tabulated using the GenomicRanges (Lawrence et al., 2013) package in R/Bioconductor. Genes expressed at 3 reads per million or more in all samples from one group were considered detected and used as input for edgeR v3.24.3 (Robinson et al., 2010) to identify DEG. P-values were FDR corrected using the Benjamin-Hochberg method (Y, 1995) with genes of an FDR <0.05 and a FC >1 or <- 1 log(2) considered significant. Expression data was normalized to reads per kilobase per million mapped reads (RPKM). Samples included either four (D14 timepoint) or six (D7 timepoint) biological replicates per B cell subset. Principal component analysis was performed using the vegan package v2.5.6. All RNA-seq data is available from the GEO database under the accession GSE163989. See also Table S2.

### ATAC-seq preparation and analysis

B cell subsets were enriched from PBMCs (EasySep^TM^ Human B Cell Negative Selection Kit), sort-purified (Table S12), re-suspended in 25 μl of tagmentation reaction buffer (2.5 μl Tn5, 1xTagment DNA Buffer, 0.02% Digitonin, 0.01% Tween-20) and incubated for 1hr at 37°C. Cells were lysed with 25 μl 2x Lysis Buffer (300 mM NaCl, 100 mL EDTA, 0.6% SDS, 1.6 μg Proteinase-K) for 30 min at 40°C. Low molecular weight DNA was purified by size-selection with SPRI-beads (Agencourt), and PCR amplified using Nextera primers with 2x HiFi Polymerase Master Mix (KAPA Biosystems). Amplified, low molecular weight DNA was isolated using SPRI-bead size selection. Libraries were quality assessed and sequenced using a 75bp paired end run on a NextSeq500 at the UAB Heflin Genomics Center. Raw sequencing reads were mapped to the hg38 version of the human genome using Bowtie (v1.1.1) (Langmead, 2010) with the default settings. Duplicate reads were annotated using the Picard Tools MarkDuplicates function (http://broadinstitute.github.io/picard/) and eliminated from downstream analysis. Enriched peaks were identified using MACS2 v2.1.0.2014061) with the default settings (Zhang et al., 2008). Genomic and motif annotations were determined for ATAC-seq peaks using the HOMER (Heinz et al., 2010) (v4.10) annotatePeaks.pl script. Read counts for all peaks were annotated for each sample from the bam file using the GenomicRanges (Lawrence et al., 2013) R/Bioconductor package and normalized to Reads Per Peak Per Million (RPPM) (Scharer et al., 2016). Differential accessible regions/peaks (DAR) were determined using edgeR v3.24.3(Robinson et al., 2010) and P-values were FDR corrected using the Benjamin-Hochberg method (Y, 1995). Peaks with a >1 or <- 1 log(2) FC and FDR < 0.05 between comparisons were termed significant. IRF4 target accessibility was computed by taking the mean peak accessibility for all peaks that mapped to genes derived from the Shaffer IRF4_Up in PCs dataset (Shaffer et al., 2008). All ATAC-seq data is available from the NCBI Gene Expression Omnibus (GEO) database under the accession GSE163989. See also Table S3.

### Gene set enrichment analyses

Detected genes in the RNA-seq datasets were ranked by multiplying the -log_10_ of the P-value from edgeR by the sign of the FC and then used as input in the GSEA (Subramanian et al., 2005) PreRanked analysis program (http://software.broadinstitute.org/gsea/index.jsp). Gene lists evaluated by GSEA include mSigDB curated Gene Ontogeny (GO) terms (Liberzon et al., 2011) version 7.0, collection C5) and selected gene sets provided in Table S4.

### Identification of signaling and metabolism transcription modules in effector memory B cells

GSEA was performed comparing the ranked gene list of D7 post-vaccine IgD^neg^ H1^+^ T-bet^hi^ and T-bet^lo^ memory B cell subsets to 3931 GO terms (mSigDB version 7.0, collection C5). Enriched gene sets with an FDR q-value < 0.01 were identified (See Table S10). A distance matrix was constructed based on 1 - Jaccard index of the constituent leading-edge genes of pairs of these enriched terms and was used for classical multidimensional scaling to visualize the terms in three-dimensional space. Terms were grouped in clusters (k-means clustering with n=9 as specified by the gap statistic (Tibshirani R, 2001)) using Matlab’s evalcluster function. Leading-edge genes from the GO lists present in each cluster were compared and 11 prototypic gene lists with the most similarity to other gene lists (i.e. largest number of shared leading-edge genes) within the cluster were selected (Table S10) for further analysis. The known properties of the proteins encoded by the leading-edge genes from each of the 11 prototypic GO lists were manually curated and the genes were grouped into 14 modules (Table S11) based on the reported functional and metabolic properties of these genes. The module assignments were independently validated using publicly available data sets for the relevant metabolic pathways.

### T-bet Motif and ChIP-seq Analysis

T-bet motifs in accessible regions were annotated using HOMER and annotatePeaks.pl [DAR peak file] hg38 -size given -noann -m tbx21.motif -mbed [tbx21.motifs.bed]. The resulting bed file was used as input to calculate the RPPM normalized accessibility in the 50bp surrounding each motif for the indicated samples. T-bet ChIP-seq data from GM12878 cells was downloaded from the GEO database under accession GSE92020 (Consortium, 2012). The overlap of chromatin accessibility at T-bet peaks was calculated using the GSE92020_ENCFF971VHK_optimal_idr_thresholded_peaks_GRCh38.bed file as described above.

### PageRank (PR) Analysis

PR analysis was performed using the D7 or D14 RNA-seq and ATAC-seq data sets from each subset as previously described (Yu et al., 2017). TFs with a PR statistic greater than 0.001 in any cell type were considered for downstream analysis. The log2 FC (log2FC) in PR statistic between subsets was calculated for each TF. See also Table S6.

### Ingenuity Pathway Analysis (IPA)

Enriched pathways were identified using IPA (QIAGEN Inc., Redwood City CA, https://www.qiagenbioinformatics.com/products/ingenuity-pathway-analysis) using a gene list derived from T-bet motif containing DAR that were significantly increased (FDR corrected *P* < 0.05, FC > 1 Log(2)) in D7 HA^+^ T-bet^hi^ relative to T-bet^lo^ memory cells (Table S7).

### B cell *in vitro* stimulation

Tonsil B cell subsets were sort-purified as described in Table S12 and stained for 10 min at 37°C with PBS diluted CellTrace^TM^ Violet (Molecular Probes, Thermofisher). Labeled cells (0.5x10^6^ cells/mL) were cultured for 2-3 days with 5 μg/mL R848 (InvivoGen), 50 U/mL IL-2 (Peprotech), 50 ng/mL IL-21 (Peprotech) and 20 ng/ml IFNγ (R&D) and analyzed.

### HA ELISAs

Plasma from vaccinated samples were serially diluted on EIA/RIA ELISA plates (Costar) that were previously coated with recombinant HA. HA-specific IgG Abs from the samples were detected using HRP- conjugated anti-human IgG secondary Abs (Jackson ImmunoResearch) and were developed using ABTS with acid stop. Absorbance was measured at 415nm using a SpectraMaxM2 (Molecular Devices). All samples were tested against the same reference standard and 50% endpoint titers were determined.

### ELISPOT

B cells from *in vitro* cultures were counted, serially diluted onto anti-IgG (Jackson ImmunoResearch) coated ELISPOT plates (Millipore) and incubated for 6 hr. Bound Ab was detected with AP-conjugated anti-human IgG (Jackson ImmunoResearch) and AP substrate (Moss, Inc). ELISPOTs were visualized and captured using a CTL ELISPOT reader for direct import into CANVAS visualization software with subsequent equivalent and proportional image reduction.

### Metabolism assays

Sort-purified B cell subsets (Table S12) were incubated for 15 min at 37°C with 2ng/mL of Mitrotracker Green (Invitrogen), washed in PBS and immediately analyzed or were fixed in 10% neutral buffered formalin (Sigma), permeabilized in 0.1% IGEPAL (Sigma) and stained with anti-pS6 Ab (Cell Signaling Technology). In other experiments cell, tonsil B cells were surface stained, washed and then incubated with either 2 ng/mL of H_2_DFCDA (Invitrogen) for 20 min at 4°C or with 5 ng/mL of BODIPY 500/510 (Invitrogen). Cells were washed with PBS and immediately analyzed.

### ImageStream

CD19^+^ cells were enriched (EasySep^TM^ human B cell negative selection kit) from HD PBMCs collected D7 post-IIV. Cells were stained with APC-conjugated H1 tetramer and Abs for IgD, CD38 and CD19 for 20 min at 4°C and then fixed 10% neutral buffered formalin (Sigma). Cells were permeabilized with 0.1% IGEPAL (Sigma) in the presence of PE-conjugated H1 tetramer and anti-T-bet Ab. Cells were analyzed using ImageStreamXMarkII (Luminex) and IDEAS software. Cells were gated on in-focus singlets. The intensity mask feature was used to acquire the intensity of extracelluar H1 tetramer and intracellular H1 tetramer staining in various B cell subsets. Single cell images were exported from IDEAS software as TIFF files with direct import into CANVAS for equivalent and proportional reduction. Final images are presented as 600 dpi.

### Statistical Analyses

Comparisons between two groups were performed with the Wilcoxson matched pairs signed rank test for non-normally distributed variables. Two-tailed t-testing was used to compare mean RPPM values for cumulative genes in the Shaffer UP in IRF4 gene set and for T-bet ChIP-seq and T-bet motif comparisons. The one-way ANOVA test was used to compare mean values of three or more groups and either the Dunn’s or Tukey’s multiple comparisons test was used to compare medians. Strength and direction of association between two variables measures was performed using the D’Agostino-Pearson normality test followed by or Spearman correlation test. Data were considered significant when *P* ≤ 0.05. GraphPad Prism version 7.0a or 8.0 software (GraphPad) was used for analysis.

### Other bioinformatics analyses and visualizations

Hierarchical clustering, PCA, heatmap plots and other visualizations were created using Matlab (R2020a, The Mathworks Inc., Natick MA) or R v3.5.2.

### Data and Code Availability

All RNA and ATAC-seq sequencing data is available from the NCBI Gene Expression Omnibus (GEO) under the series GSE163989. Select code is available from https://github.com/cdschar/ and specific analysis routines are available upon request. Ig-Rep-seq processed data files are also available upon request.

---

**Figure S1.**
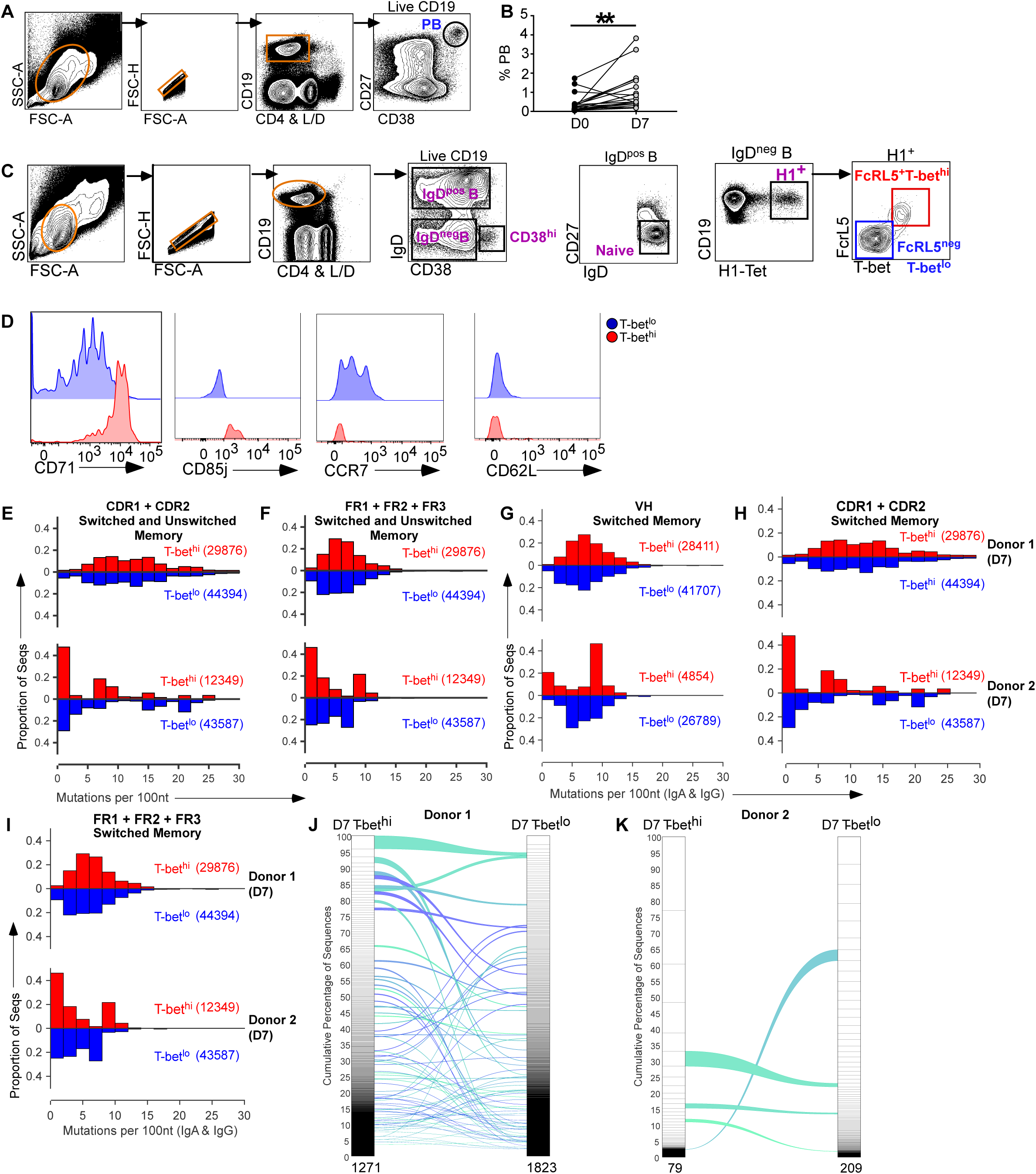
Supporting Information for Figure 1. (**A-C**) B lineage subsets in peripheral blood of HD (n=19) immunized with IIV (2015). Representative flow cytometry plots defining the CD38^hi^CD27^hi^ PB population (**A**), IgD^+^CD27^neg^, naïve B (**C**), IgD^neg^ H1-specific B (**C**) and the IgD^neg^ H1-specific FcRL5^+^T-bet^hi^ (**C**) and FcRL5^neg^T-bet^lo^ (**C**) cells. Enumerated PBs on D0 and D7 post-IIV (**B**) shown as frequency of PBs within the total live CD19^+^ B cell subset for each HD. **p<0.001 paired Student’s t-test. (**D**) Histograms showing expression of CD71, CD85j, CCR7, and CD62L by the H1^+^ T-bet^hi^ (red) and H1^+^ T-bet^lo^ (blue) IgD^neg^ B cells. (**E-K**) IgH(V_H_) BCR repertoire analysis of sort-purified (gating in Fig. S1A,C) H3^+^ IgD^neg^ T-bet^hi^ and T-bet^lo^ B cells (n=2 HD) on D7 post-IIV (2016). V_H_ mutations represented as the frequency of mutations per 100 nt in the CDR1+CDR2 (**E**) or FR1+FR2+FR3 (**F**) domain of the sorted B cell populations. Data also shown as the frequency of mutations per 100 nt in the V_H_ (**G**), CDR1+CDR2 (**H**) or FR1+FR2+FR3 (**I**) domains of isotype-switched (IgA^+^ or IgG^+^) B cell populations. Number of sequences in each population is indicated. Shared lineages, identified between D7 H3^+^ IgD^neg^ T-bet^hi^ and T-bet^lo^ cells, are represented as alluvial plots (**K**) with the individual lineages ranked by size in each subset. Total number of lineages indicated at the bottom of each population bar. See Table S1E-F for BCR repertoire data.

**Figure S2.**
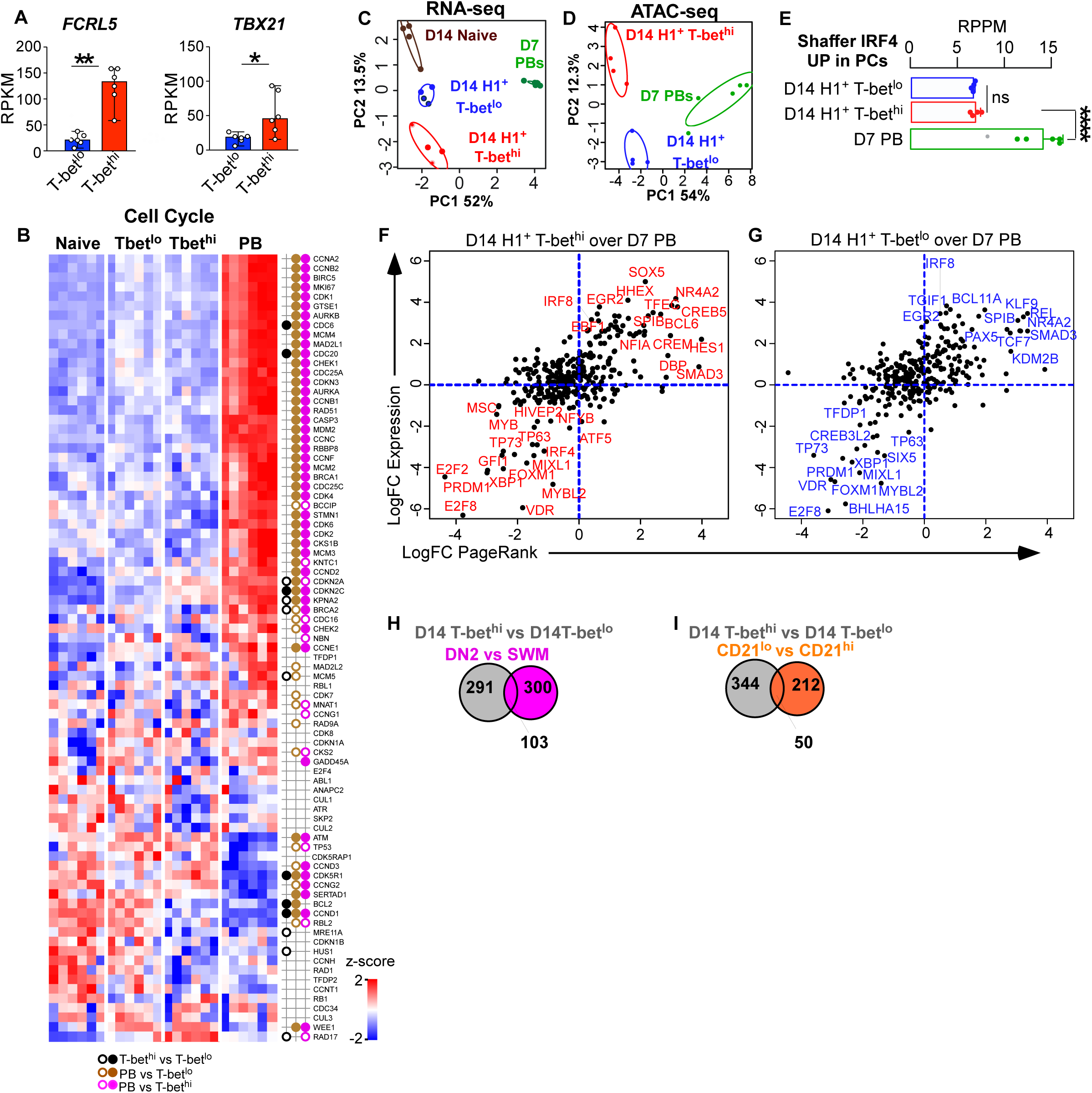
Supporting information for Figure 2. (A) RNA expression levels (in RPKM) for *TBX21* and *FCRL5* in D7 post-IIV H1^+^ IgD^neg^ T-bet^hi^ and T-bet^lo^ cells. (**B-G**) RNA-seq and ATAC-seq analyses was performed on circulating PB (green), naïve B cells (brown), H1^+^ IgD^neg^ T-bet^lo^ (FCRL5^neg^, blue) cells and H1^+^ IgD^neg^ T-bet^hi^ B cells (FCRL5^+^, red) that were sort-purified (Fig. S1A, C for gating) from HD on D7 (**B**, n=6) or D14 (**C-G**, n=4) post-IIV. (B) RNA expression levels of cell cycle genes in D7 subsets (Table S4) represented as a heat map of z-score of log gene expression. Black circles denote comparisons between H1^+^ T-bet^hi^ vs H1^+^ T-bet^lo^, brown circles denote comparisons between PB vs H1^+^ T-bet^lo^ cells and pink circles denote comparisons between PBs vs H1^+^ T-bet^hi^ cells. Open circles indicate genes with FDR q< 0.05 and filled circles indicate genes with FDR<0.05 and >2 fold-changes up or down in expression levels. (**C-D**) PCA for the D14 RNA-seq (**C**) and ATAC-seq (**D**) data sets from each B cell subset. (**E**) Chromatin accessibility surrounding IRF4 target genes in human plasma cells (Table S4, (Shaffer et al., 2008)) as assessed in D14 ATAC-seq data. Data, reported as RPPM, represent mean peak accessibility for all peaks mapping to genes directly regulated by IRF4 in plasma cells. (**F-G**) TFs identified by PR as regulators of D7 PBs, D14 H1^+^ IgD^neg^ T-bet^hi^ and D14 H1^+^ IgD^neg^ T-bet^lo^ gene networks (Table S6). Comparison between the T-bet^hi^ memory B cells over PB (**F**) or T-bet^lo^ memory B cells over PBs (**G**) with individual TFs indicated. (**H-I**) Venn diagrams showing overlap between the 394 DEGs identified in D14 H1^+^ IgD^neg^ T-bet^hi^ vs T-bet^lo^ memory B cells and (**H**) the 403 DEG identified in DN2 vs SWM B cells in SLE patients (Table S5C, (Jenks et al., 2018)) or (**I**) between the 262 DEG identified in CD19^+^ CD38^lo/med^ CD27^+^CD21^lo^ vs CD21^hi^ B cells D14 post-IIV (Table S5D, (Lau et al., 2017)). Statistical analyses were performed using Wilcoxson matched pairs signed rank test (**A)** or two tailed t-testing (**E**). *p< 0.05, **, p<0.01, *** p<0.001, **** p <0.0001 ns= non-significant. p_nom_ values for GSEA analyses are indicated.

**Figure S3.**
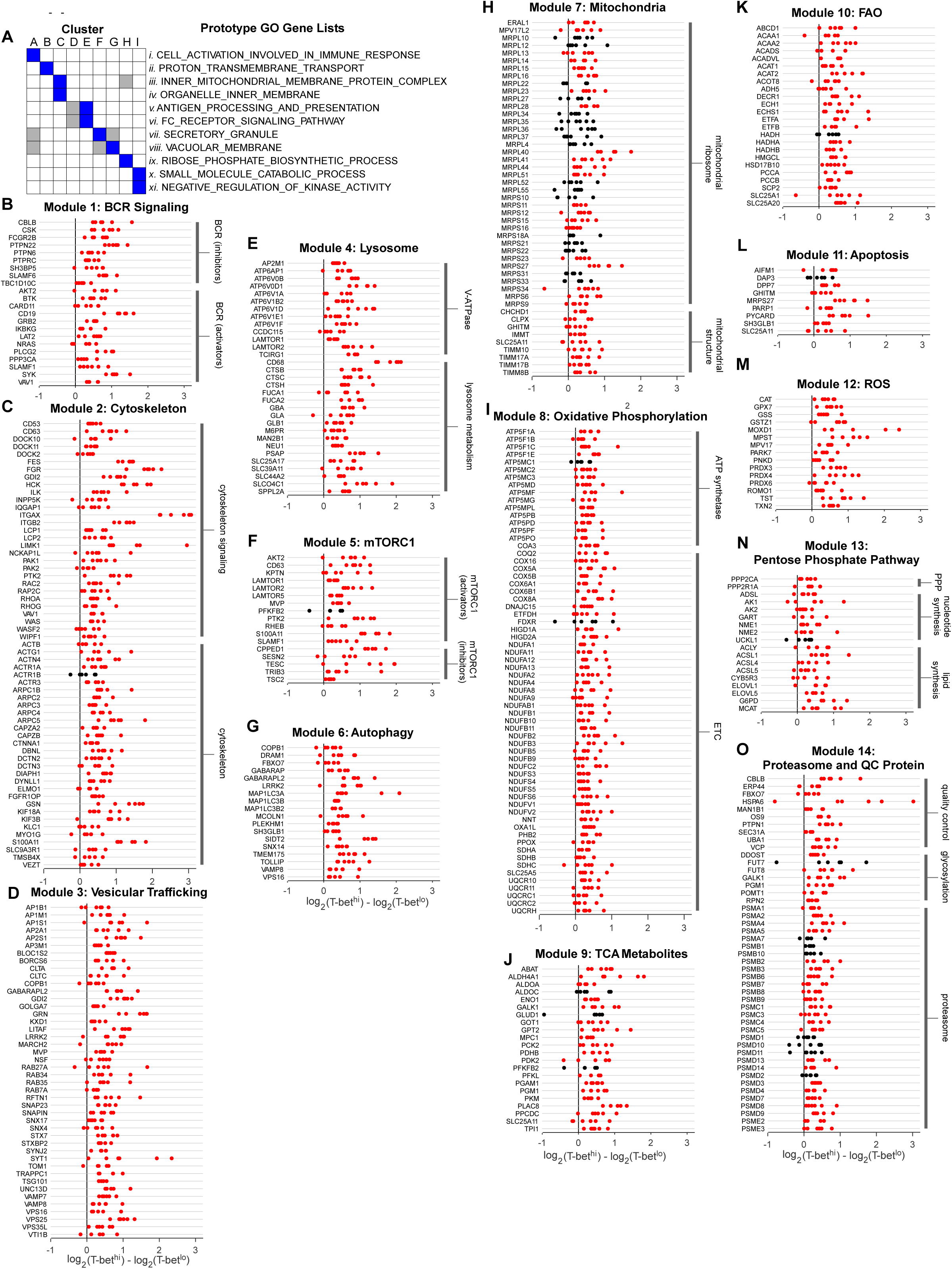
Supporting information for Figure 5. **(A)** GSEA comparing the RNA-seq ranked gene list from D7 H1^+^ IgD^neg^ T-bet^hi^ and T-bet^lo^ B cells to 3931 GO gene lists was performed. GO gene lists (n = 185) significantly enriched (FDR q-value < 0.01,) for expression in T-bet^hi^ relative to the T-bet^lo^ memory B cells were analyzed by multidimensional scaling and clustering (9 clusters (referred to as A-I), Table S10; see methods). Eleven prototypic GO gene lists (referred to as *i-xi*) representing the 9 GO gene set clusters were selected (Table S10). Relationship between the 11 Prototype GO lists (*i-xi*) and the 9 clusters (A-I) is shown. Some GO lists were associated with more than 1 cluster (blue, primary cluster and grey, secondary cluster) and more than 1 prototypic GO list was identified for some clusters. (**B-O**). The leading-edge genes from the 11 prototypic GO gene lists were assigned to 14 signaling and metabolic modules (**B-O,** Table S11). Expression of the genes assigned to the 14 modules in the D7 H1^+^ IgD^neg^ T-bet^hi^ and T-bet^lo^ memory B cells is shown with red dots indicating genes from individual RNA-seq samples with FDR q<0.05 in the T-bet^hi^ over T-bet^lo^ comparison and black dots indicating genes with FDR q>0.05. Vertical black bars indicate functional subdivisions within each module.

**Figure S4.**
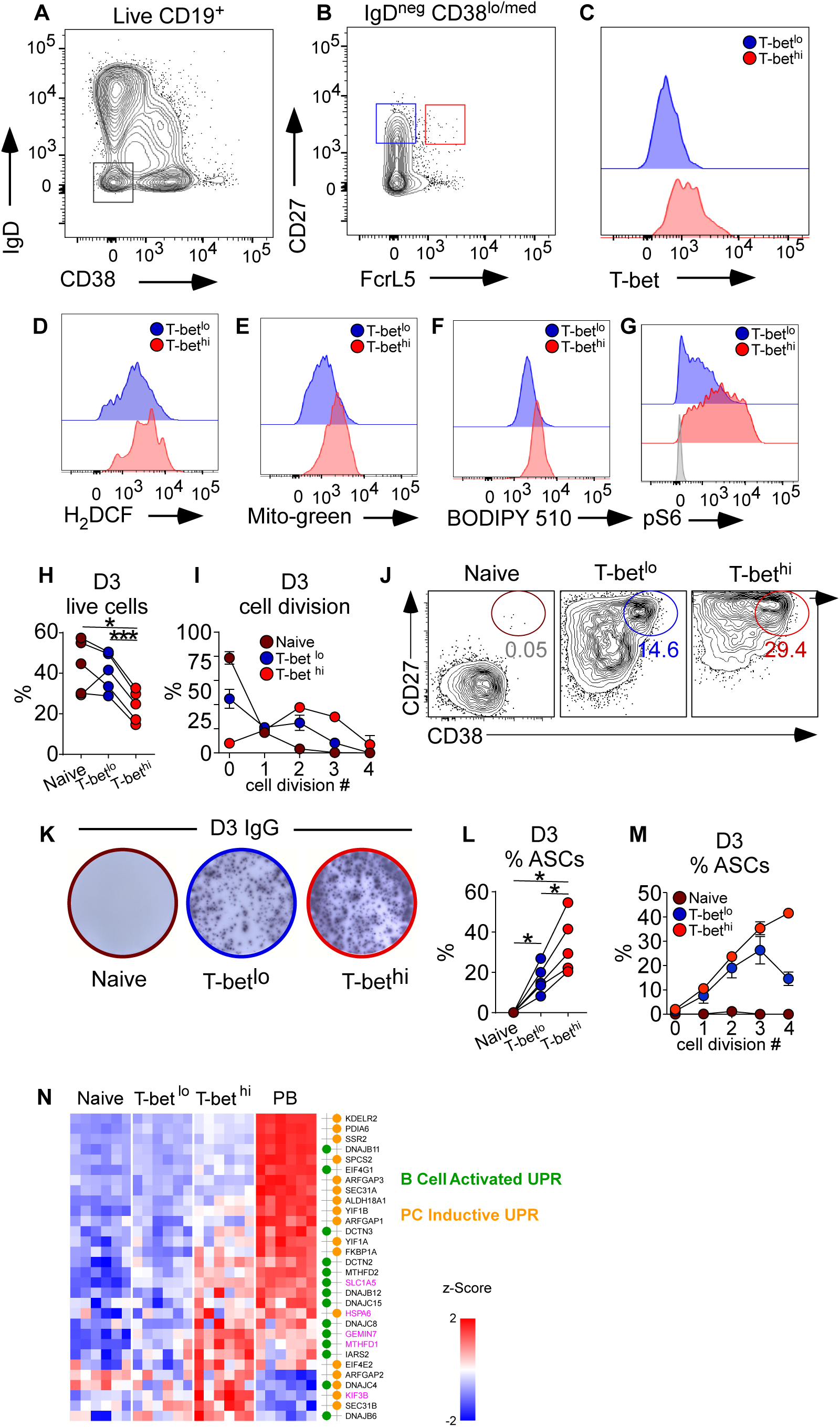
Supporting Information for Figure 6. (**A-C**) Gating strategy to identify T-bet^hi^ and T-bet^lo^ IgD^neg^CD27^+^ memory B cells in tonsil. Non-GC, Ag-experienced CD38^lo^IgD^neg^ CD19^+^ tonsil B cells (**A**) were analyzed for expression of CD27, FcRL5 and T-bet (**B-C**). The FcRL5^+^ IgD^neg^CD27^+^ memory B cells (**B**, red gate) expressed high levels of T-bet (**C**) while the FcRL5^neg^ IgD^neg^CD27^+^ memory B cells (**B**, blue gate) expressed uniformly low levels of T-bet (**C**). (**D-G**) Metabolic analyses of tonsil-derived matched T-bet^hi^ (FcRL5^+^, red) and T-bet^lo^ (FcRL5^neg^, blue) IgD^neg^CD27^+^ memory B cells showing representative histograms of T-bet^hi^ and T-bet^lo^ cells from the same donor. (D) ROS levels assessed by flow cytometry following staining with 2, 7 – dichlorohydrofluorescein diacetate (H_2_DCF). (E) Mitochondrial mass as measured by staining with Mitotracker Green. (F) Membrane lipid composition analyzed by staining with the fatty acid analog BODIPY510. (G) mTORC1 activation as assessed by expression of phosphorylated S6 (pS6) kinase. Gray histogram indicates isotype control. (**H-M**) Donor matched naïve (brown), T-bet^hi^ (FcrL5^+^, red) or T-bet^lo^ (FcrL5^neg^, blue) IgD^neg^CD27^+^CD38^lo/med^ memory B cells were sort-purified (see Fig. S4A-C for sorting strategy), labeled with CTV, stimulated with R848, IL-21, IL-2 and IFN-γ and assessed on D3. **(H-I)** Cell survival (**H**) and frequency of cells in each cell division (**I**) determined by flow cytometry. (**J-M**) ASCs in cultures enumerated by flow cytometry (**J**) and ELISPOT **(K)**. Frequencies of total CD27^hi^CD38^hi^ ASCs (**L**) from each matched culture and percentage of ASCs in each cell division (**M**) are shown. (**N**) RNA expression levels of genes upregulated by the mTORC1 dependent early UPR in mouse follicular B cells (Gaudette et al., 2020) (Table S4) activated with CpG (B cell activation UPR, green circles) or CpG, IL-4 and IL-5 (PC inductive UPR, orange circles). Data shown as heat map of z-score of log gene expression in D7 naïve B cells, T-bet^lo^ and T-bet^hi^ memory B cells and PBs.

**Figure S5.**
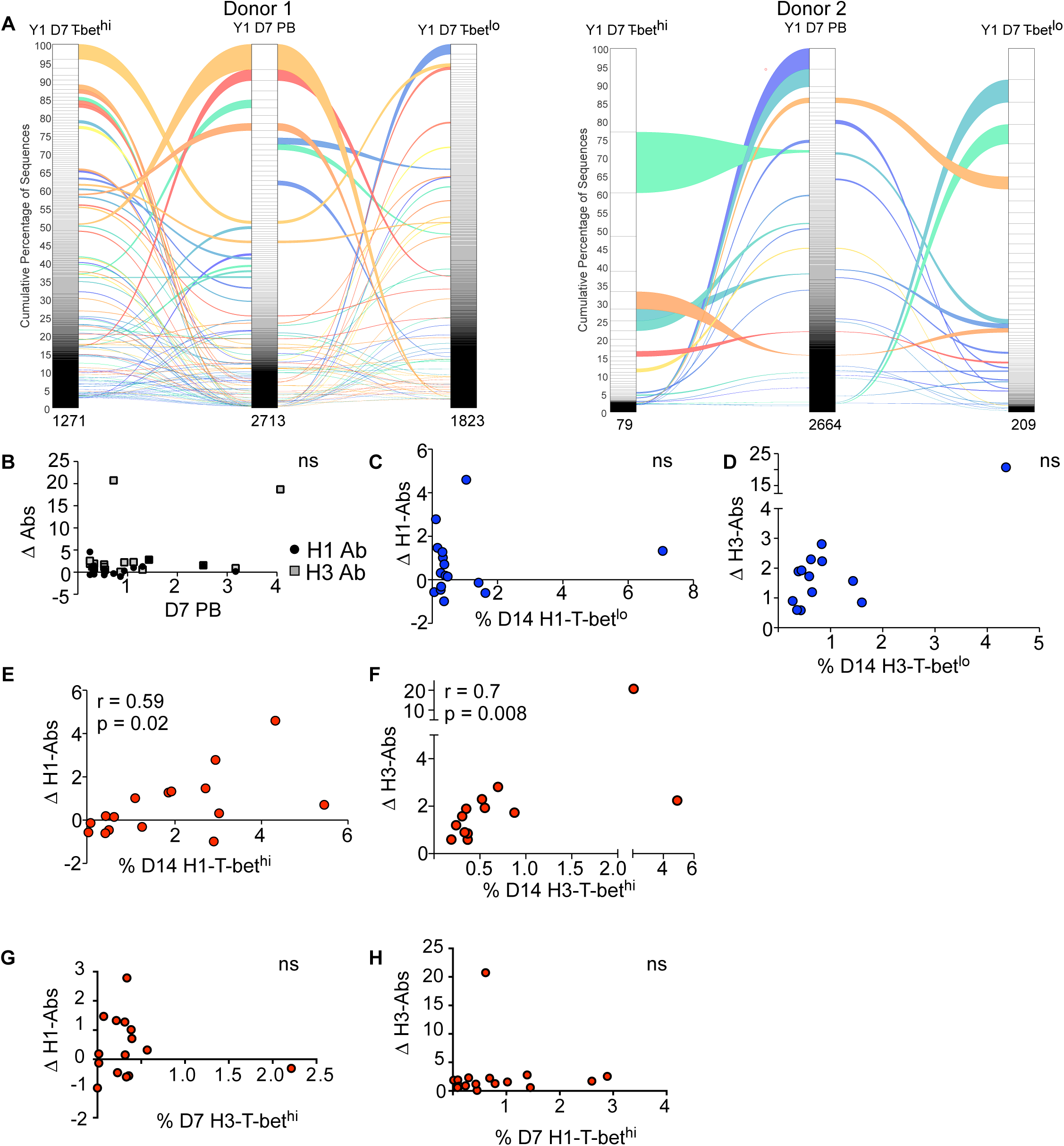
Supporting information for Figure 7. (A) IgH(V_H_) BCR repertoire analysis to identify shared lineages between PBs and H1^+^ IgD^neg^ T-bet^hi^ and T-bet^lo^ B cells isolated from HD (n=2) on D7 post-IIV (2016). Data showing connectivity of shared lineages represented as alluvial plots with the individual lineages ranked by size in each subset. Ribbons identify lineages shared with the PB population. Lineages that are shared between PB and one H1^+^ IgD^neg^ population are colorized on green-blue scale and lineages shared by PB and both H1^+^ IgD^neg^ populations are colorized on yellow-red scale. Total number of lineages indicated at the bottom of each population bar. See Table S1E-F for BCR repertoire data. (**B-H**) Assessment of vaccine-specific D7 and D14 memory PB and B cell responses (measured as frequency of population in blood) and vaccine-specific Ab responses (measured as FC) in titers between D0 and D120) after IIV (2015) in HD (n=19). (B) Correlation between the FC in H1- (circles) and H3- (gray squares) specific IgG titers (Y axis) and the frequency of the D7 PBs expressed within the CD19^+^ B cell subset (X axis). (**C-D**) Correlation between FC in H1- (**C**) and H3- (**D**) specific IgG titers (Y axis) and the frequency of the D14 H1- (**C**) and H3- (**D**) specific T-bet^lo^ cells within the IgD^neg^ B subset (X axis). (**E-F**) Correlation between the FC in H1- (**E**) and H3- (**F**) specific IgG titers (Y axis) and the frequency of the D14 H1- (**E**) and H3- (**F**) specific T-bet^hi^ cells within the IgD^neg^ B subset (X axis). (**G-H**) Correlation between the FC in H1- (**G**) or H3- (**H**) specific IgG titers (Y axis) and the frequencies of the *reciprocal* D7 H3- (**G**) or H1- (**H**) specific T-bet^hi^ IgD^neg^ B cells within the IgD^neg^ B cell subset (X axis). ns = no significant correlation in indicated comparisons. Spearman correlation (r) and significance (p) between indicated correlations are shown.

**Supplemental Table S1. Avidity measurements of recombinant Abs generated from single sort-purified H1^+^ IgD^neg^ B cells and BCR repertoire data.** Related to Figures 1, 3, 7, S1 and S5. Tab 1A contains BCR reactivity data (reported as gMFI) for 270 recombinant H1-specific Abs (rAb) generated from one IIV-immunized HD. Tab 1B provides statistical analyses associated with rAb reactivity. Tab 1C-F delineates BCR repertoire data acquired from (n=3) IIV immunized HD.

**Supplemental Table S2. RNA-seq data set from sort-purified B cell subsets D7 and D14 days after influenza vaccination.** Related to Figures 2, 3, 4, 5, 6, S2, S3 and S4. RNA-seq analysis of sort-purified B cell subsets isolated from the peripheral blood of HD on D7 (n=6 donors) and D14 (n=4 donors) post-IIV. Data reported as RPKM values for each gene. Log(2)FC, p values and FDR values provided for each B cell subset pairwise comparison.

**Supplemental Table S3. ATAC-seq dataset from sort-purified B cell subsets D7 and D14 after influenza vaccination.** Related to Figures 2, 3, 4, and S2. ATAC-seq analysis of sort-purified B cell subsets isolated from the peripheral blood of HD on D7 (n=5 donors) and D14 (n=4 donors) post-IIV. Data reported as RPPM values for each DAR. Log(2)FC, p values and FDR values provided for each B cell subset pairwise comparison.

**Supplemental Table S4. Gene sets used in GSEA.** Related to Figures 2, 3, 5, and 6. Description of all gene sets used for GSEA including the source of the gene sets and the list of genes in each set.

**Supplemental Table S5. Comparison of DEG between different B cell subsets.** Related to Figure 2 and S2. DEG from D7 and D14 RNA-seq datasets (derived from Table S2) and published datasets in Jenks et al (Jenks et al., 2018) and Lau et al (Lau et al., 2017) (See Table S4 for lists) are displayed. Genes are reported as significant (1) or not significant (0) in the comparator gene set. Tabs A-B include D7 comparisons and Tabs C- D include D14 comparisons.

**Supplemental Table S6. PR identification of putative transcriptional regulators of B cell subsets on D7 and D14 after vaccination.** Related to Figures 2, 3 and S2. D7 and D14 RNA-seq (Table S2) and ATAC-seq (Table S3) data derived from D7 PB and D7 and D14 H1^+^ IgD^neg^ T-bet^hi^ and T-bet^lo^ memory B cell subsets were analyzed using the PR algorithm to identify TFs predicted to regulate the transcriptional networks in each B cell subset. Data are reported as PR scores for TFs in each subset and as FClog(2) in PR scores between different B cell subsets.

**Supplemental Table S7. Identification of T-bet motif-containing DAR.** Related to Figure 4. Identification of 468 DAR (derived from Table S3) that were (a) significantly up in T-bet^hi^ memory B cells over T-bet^lo^ memory B cells and (b) contained a consensus T-bet binding motif. The 468 DAR meeting these requirements mapped to 402 unique genes. FC and FDR values for gene expression of these 402 genes are provided. FC and FDR values for **ALL** DAR (n=553) assigned to these 402 genes and the number of T-bet binding motifs associated with each DAR are reported.

**Supplemental Table S8. Pathway analysis of genes with exhibiting epigenetic changes in T-bet^hi^ memory B cells.** Related to Figure 4. Genes (n=402), which were identified in Table S7 as linked to at least one T-bet motif containing DAR that exhibited increased chromatin accessibility in D7 T-bet^hi^ memory B cells, were evaluated using IPA. Top IPA-predicted pathways are reported.

**Supplemental Table S9. Epigenetic changes in transcriptional regulators that are differentially expressed in T-bet^hi^ and T-bet^lo^ memory B cells.** Related to Figure 4. Intersection of the DEG (Table S2) and DAR (Table S3) between T-bet^hi^ and T-bet^lo^ memory B cells (111 unique genes assigned to 154 DAR). DAR with consensus T-bet binding motifs or containing known ChIP-seq assigned T-bet binding sites (Consortium, 2012) are indicated. Gene annotation and putative functions are provided.

**Supplemental Table S10. GSEA of RNA-seq data from D7 H1^+^ IgD^neg^ T-bet^hi^ versus T-bet^lo^ B cell subsets against representative GO gene sets.** Related to Figure 5 and S3. GSEA comparison of the ranked gene list of D7 post-vaccine H1^+^ IgD^neg^ T-bet^hi^ and T-bet^lo^ memory B cell subsets to 3931 GO terms. Identification of 185 positively enriched gene sets with an FDR q-value < 0.01 (Tab 1). Nine clusters of gene sets (labeled Tabs A-I) with overlapping leading-edge genes were identified. Leading edge-genes for each gene list in each cluster are reported as present (1) or absent (0). Prototypic gene lists for each cluster are highlighted in yellow.

**Supplemental Table S11. Functional Gene Modules.** Related to Figure 5 and S3. Functional descriptions for proteins encoded by leading-edge genes from 11 prototypic gene sets (Table S10) were curated and each gene was assigned to a functional module (n=14 modules). Table shows gene set origin of each leading-edge gene and assignment of leading-edge genes into the different modules.

**Supplemental Table S12. Sort Panels**. Related to all Figures. Flow panels for various flow sort purifications of B cell subsets outlined.

**Supplemental Table S13. Primers for rAb generation.** Related to Figure 1. Primers used to amplify the heavy and light chain of single sorted cells for recombinant monoclonal Ab generation.

## References

Alamyar, E., Duroux, P., Lefranc, M.P., and Giudicelli, V. (2012). IMGT((R)) tools for the nucleotide analysis of immunoglobulin (IG) and T cell receptor (TR) V-(D)-J repertoires, polymorphisms, and IG mutations: IMGT/V-QUEST and IMGT/HighV-QUEST for NGS. Methods Mol Biol 882, 569–604.

Allie, S.R., Bradley, J.E., Mudunuru, U., Schultz, M.D., Graf, B.A., Lund, F.E., and Randall, T.D. (2019). The establishment of resident memory B cells in the lung requires local antigen encounter. Nat Immunol 20, 97–108.

Andrews, S.F., Chambers, M.J., Schramm, C.A., Plyler, J., Raab, J.E., Kanekiyo, M., Gillespie, R.A., Ransier, A., Darko, S., Hu, J., et al. (2019). Activation Dynamics and Immunoglobulin Evolution of Pre-existing and Newly Generated Human Memory B cell Responses to Influenza Hemagglutinin. Immunity 51, 398–410 e395.

Andrews, S.F., Huang, Y., Kaur, K., Popova, L.I., Ho, I.Y., Pauli, N.T., Henry Dunand, C.J., Taylor, W.M., Lim, S., Huang, M., et al. (2015). Immune history profoundly affects broadly protective B cell responses to influenza. Sci Transl Med 7, 316ra192.

Benhamron, S., Pattanayak, S.P., Berger, M., and Tirosh, B. (2015). mTOR activation promotes plasma cell differentiation and bypasses XBP-1 for immunoglobulin secretion. Mol Cell Biol 35, 153–166.

Bentebibel, S.E., Lopez, S., Obermoser, G., Schmitt, N., Mueller, C., Harrod, C., Flano, E., Mejias, A., Albrecht, R.A., Blankenship, D., et al. (2013). Induction of ICOS+CXCR3+CXCR5+ TH cells correlates with antibody responses to influenza vaccination. Sci Transl Med 5, 176ra132.

Bertolotti, M., Sitia, R., and Rubartelli, A. (2012). On the redox control of B lymphocyte differentiation and function. Antioxid Redox Signal 16, 1139–1149.

Boonyaratanakornkit, J., and Taylor, J.J. (2019). Techniques to Study Antigen-Specific B Cell Responses. Front Immunol 10, 1694.

Buck, M.D., Sowell, R.T., Kaech, S.M., and Pearce, E.L. (2017). Metabolic Instruction of Immunity. Cell 169, 570–586.

Consortium, E.P. (2012). An integrated encyclopedia of DNA elements in the human genome. Nature 489, 57–74.

Dobin, A., Davis, C.A., Schlesinger, F., Drenkow, J., Zaleski, C., Jha, S., Batut, P., Chaisson, M., and Gingeras, T.R. (2013). STAR: ultrafast universal RNA-seq aligner. Bioinformatics 29, 15–21.

Doherty, E., and Perl, A. (2017). Measurement of Mitochondrial Mass by Flow Cytometry during Oxidative Stress. React Oxyg Species (Apex) 4, 275–283.

Ellebedy, A.H., Jackson, K.J., Kissick, H.T., Nakaya, H.I., Davis, C.W., Roskin, K.M., McElroy, A.K., Oshansky, C.M., Elbein, R., Thomas, S., et al. (2016). Defining antigen-specific plasmablast and memory B cell subsets in human blood after viral infection or vaccination. Nat Immunol 17, 1226–1234.

Elsner, R.A., and Shlomchik, M.J. (2020). Germinal Center and Extrafollicular B Cell Responses in Vaccination, Immunity, and Autoimmunity. Immunity 53, 1136–1150.

Gaudette, B.T., Jones, D.D., Bortnick, A., Argon, Y., and Allman, D. (2020). mTORC1 coordinates an immediate unfolded protein response-related transcriptome in activated B cells preceding antibody secretion. Nat Commun 11, 723.

Gunn, K.E., and Brewer, J.W. (2006). Evidence that marginal zone B cells possess an enhanced secretory apparatus and exhibit superior secretory activity. J Immunol 177, 3791–3798.

Heinz, S., Benner, C., Spann, N., Bertolino, E., Lin, Y.C., Laslo, P., Cheng, J.X., Murre, C., Singh, H., and Glass, C.K. (2010). Simple combinations of lineage-determining transcription factors prime cis-regulatory elements required for macrophage and B cell identities. Mol Cell 38, 576–589.

Hipp, N., Symington, H., Pastoret, C., Caron, G., Monvoisin, C., Tarte, K., Fest, T., and Delaloy, C. (2017). IL-2 imprints human naive B cell fate towards plasma cell through ERK/ELK1-mediated BACH2 repression. Nat Commun 8, 1443.

Igarashi, K., Ochiai, K., Itoh-Nakadai, A., and Muto, A. (2014). Orchestration of plasma cell differentiation by Bach2 and its gene regulatory network. Immunol Rev 261, 116–125.

Iwakoshi, N.N., Lee, A.H., Vallabhajosyula, P., Otipoby, K.L., Rajewsky, K., and Glimcher, L.H. (2003). Plasma cell differentiation and the unfolded protein response intersect at the transcription factor XBP-1. Nat Immunol 4, 321–329.

Jenks, S.A., Cashman, K.S., Zumaquero, E., Marigorta, U.M., Patel, A.V., Wang, X., Tomar, D., Woodruff, M.C., Simon, Z., Bugrovsky, R., et al. (2018). Distinct Effector B Cells Induced by Unregulated Toll-like Receptor 7 Contribute to Pathogenic Responses in Systemic Lupus Erythematosus. Immunity 49, 725–739 e726.

Johnson, J.L., Rosenthal, R.L., Knox, J.J., Myles, A., Naradikian, M.S., Madej, J., Kostiv, M., Rosenfeld, A.M., Meng, W., Christensen, S.R., et al. (2020). The Transcription Factor T-bet Resolves Memory B Cell Subsets with Distinct Tissue Distributions and Antibody Specificities in Mice and Humans. Immunity 52, 842–855 e846.

Kaech, S.M., Hemby, S., Kersh, E., and Ahmed, R. (2002). Molecular and functional profiling of memory CD8 T cell differentiation. Cell 111, 837–851.

Kallies, A., and Good-Jacobson, K.L. (2017). Transcription Factor T-bet Orchestrates Lineage Development and Function in the Immune System. Trends Immunol 38, 287–297.

Knox, J.J., Buggert, M., Kardava, L., Seaton, K.E., Eller, M.A., Canaday, D.H., Robb, M.L., Ostrowski, M.A., Deeks, S.G., Slifka, M.K., et al. (2017). T-bet+ B cells are induced by human viral infections and dominate the HIV gp140 response. JCI Insight 2.

Kometani, K., Nakagawa, R., Shinnakasu, R., Kaji, T., Rybouchkin, A., Moriyama, S., Furukawa, K., Koseki, H., Takemori, T., and Kurosaki, T. (2013). Repression of the transcription factor Bach2 contributes to predisposition of IgG1 memory B cells toward plasma cell differentiation. Immunity 39, 136–147.

Koutsakos, M., Wheatley, A.K., Loh, L., Clemens, E.B., Sant, S., Nussing, S., Fox, A., Chung, A.W., Laurie, K.L., Hurt, A.C., et al. (2018). Circulating TFH cells, serological memory, and tissue compartmentalization shape human influenza-specific B cell immunity. Sci Transl Med 10.

Kurachi, M., Barnitz, R.A., Yosef, N., Odorizzi, P.M., DiIorio, M.A., Lemieux, M.E., Yates, K., Godec, J., Klatt, M.G., Regev, A., et al. (2014). The transcription factor BATF operates as an essential differentiation checkpoint in early effector CD8+ T cells. Nat Immunol 15, 373–383.

Laidlaw, B.J., and Cyster, J.G. (2020). Transcriptional regulation of memory B cell differentiation. Nat Rev Immunol.

Lam, W.Y., and Bhattacharya, D. (2018). Metabolic Links between Plasma Cell Survival, Secretion, and Stress. Trends Immunol 39, 19–27.

Lam, W.Y., Jash, A., Yao, C.H., D’Souza, L., Wong, R., Nunley, R.M., Meares, G.P., Patti, G.J., and Bhattacharya, D. (2018). Metabolic and Transcriptional Modules Independently Diversify Plasma Cell Lifespan and Function. Cell Rep 24, 2479–2492 e2476.

Langmead, B. (2010). Aligning short sequencing reads with Bowtie. Curr Protoc Bioinformatics Chapter 11, Unit 11 17.

Lau, D., Lan, L.Y., Andrews, S.F., Henry, C., Rojas, K.T., Neu, K.E., Huang, M., Huang, Y., DeKosky, B., Palm, A.E., et al. (2017). Low CD21 expression defines a population of recent germinal center graduates primed for plasma cell differentiation. Sci Immunol 2.

Lawrence, M., Huber, W., Pages, H., Aboyoun, P., Carlson, M., Gentleman, R., Morgan, M.T., and Carey, V.J. (2013). Software for computing and annotating genomic ranges. PLoS Comput Biol 9, e1003118.

Lee, F.E., Halliley, J.L., Walsh, E.E., Moscatiello, A.P., Kmush, B.L., Falsey, A.R., Randall, T.D., Kaminiski, D.A., Miller, R.K., and Sanz, I. (2011). Circulating human antibody-secreting cells during vaccinations and respiratory viral infections are characterized by high specificity and lack of bystander effect. J Immunol 186, 5514–5521.

Li, H., Borrego, F., Nagata, S., and Tolnay, M. (2016). Fc Receptor-like 5 Expression Distinguishes Two Distinct Subsets of Human Circulating Tissue-like Memory B Cells. J Immunol 196, 4064–4074.

Liberzon, A., Subramanian, A., Pinchback, R., Thorvaldsdottir, H., Tamayo, P., and Mesirov, J.P. (2011). Molecular signatures database (MSigDB) 3.0. Bioinformatics 27, 1739–1740.

Lin, D., Ippolito, G.C., Zong, R.T., Bryant, J., Koslovsky, J., and Tucker, P. (2007). Bright/ARID3A contributes to chromatin accessibility of the immunoglobulin heavy chain enhancer. Mol Cancer 6, 23.

Luckey, C.J., Bhattacharya, D., Goldrath, A.W., Weissman, I.L., Benoist, C., and Mathis, D. (2006). Memory T and memory B cells share a transcriptional program of self-renewal with long-term hematopoietic stem cells. Proc Natl Acad Sci U S A 103, 3304–3309.

Ma, Y., Shimizu, Y., Mann, M.J., Jin, Y., and Hendershot, L.M. (2010). Plasma cell differentiation initiates a limited ER stress response by specifically suppressing the PERK-dependent branch of the unfolded protein response. Cell Stress Chaperones 15, 281–293.

Mesin, L., Schiepers, A., Ersching, J., Barbulescu, A., Cavazzoni, C.B., Angelini, A., Okada, T., Kurosaki, T., and Victora, G.D. (2020). Restricted Clonality and Limited Germinal Center Reentry Characterize Memory B Cell Reactivation by Boosting. Cell 180, 92–106 e111.

Nakaya, H.I., Hagan, T., Duraisingham, S.S., Lee, E.K., Kwissa, M., Rouphael, N., Frasca, D., Gersten, M., Mehta, A.K., Gaujoux, R., et al. (2015). Systems Analysis of Immunity to Influenza Vaccination across Multiple Years and in Diverse Populations Reveals Shared Molecular Signatures. Immunity 43, 1186–1198.

Oestreich, K.J., and Weinmann, A.S. (2012). T-bet employs diverse regulatory mechanisms to repress transcription. Trends Immunol 33, 78–83.

Oikonomou, C., and Hendershot, L.M. (2020). Disposing of misfolded ER proteins: A troubled substrate’s way out of the ER. Mol Cell Endocrinol 500, 110630.

Omilusik, K.D., and Goldrath, A.W. (2019). Remembering to remember: T cell memory maintenance and plasticity. Curr Opin Immunol 58, 89–97.

Pais Ferreira, D., Silva, J.G., Wyss, T., Fuertes Marraco, S.A., Scarpellino, L., Charmoy, M., Maas, R., Siddiqui, I., Tang, L., Joyce, J.A., et al. (2020). Central memory CD8(+) T cells derive from stem-like Tcf7(hi) effector cells in the absence of cytotoxic differentiation. Immunity 53, 985–1000 e1011.

Pengo, N., Scolari, M., Oliva, L., Milan, E., Mainoldi, F., Raimondi, A., Fagioli, C., Merlini, A., Mariani, E., Pasqualetto, E., et al. (2013). Plasma cells require autophagy for sustainable immunoglobulin production. Nat Immunol 14, 298–305.

Price, M.J., Patterson, D.G., Scharer, C.D., and Boss, J.M. (2018). Progressive Upregulation of Oxidative Metabolism Facilitates Plasmablast Differentiation to a T-Independent Antigen. Cell Rep 23, 3152–3159.

Raybuck, A.L., Lee, K., Cho, S.H., Li, J., Thomas, J.W., and Boothby, M.R. (2019). mTORC1 as a cell-intrinsic rheostat that shapes development, preimmune repertoire, and function of B lymphocytes. FASEB J 33, 13202–13215.

Robinson, M.D., McCarthy, D.J., and Smyth, G.K. (2010). edgeR: a Bioconductor package for differential expression analysis of digital gene expression data. Bioinformatics 26, 139–140.

Sanz, I., Wei, C., Jenks, S.A., Cashman, K.S., Tipton, C., Woodruff, M.C., Hom, J., and Lee, F.E. (2019). Challenges and Opportunities for Consistent Classification of Human B Cell and Plasma Cell Populations. Front Immunol 10, 2458.

Saxton, R.A., and Sabatini, D.M. (2017). mTOR Signaling in Growth, Metabolism, and Disease. Cell 168, 960–976.

Scharer, C.D., Barwick, B.G., Guo, M., Bally, A.P.R., and Boss, J.M. (2018). Plasma cell differentiation is controlled by multiple cell division-coupled epigenetic programs. Nat Commun 9, 1698.

Scharer, C.D., Blalock, E.L., Barwick, B.G., Haines, R.R., Wei, C., Sanz, I., and Boss, J.M. (2016). ATAC-seq on biobanked specimens defines a unique chromatin accessibility structure in naive SLE B cells. Sci Rep 6, 27030.

Scharer, C.D., Blalock, E.L., Mi, T., Barwick, B.G., Jenks, S.A., Deguchi, T., Cashman, K.S., Neary, B.E., Patterson, D.G., Hicks, S.L., et al. (2019). Epigenetic programming underpins B cell dysfunction in human SLE. Nat Immunol 20, 1071–1082.

Schmidlin, H., Diehl, S.A., Nagasawa, M., Scheeren, F.A., Schotte, R., Uittenbogaart, C.H., Spits, H., and Blom, B. (2008). Spi-B inhibits human plasma cell differentiation by repressing BLIMP1 and XBP-1 expression. Blood 112, 1804–1812.

Shaffer, A.L., Emre, N.C., Lamy, L., Ngo, V.N., Wright, G., Xiao, W., Powell, J., Dave, S., Yu, X., Zhao, H., et al. (2008). IRF4 addiction in multiple myeloma. Nature 454, 226–231.

Shaffer, A.L., Shapiro-Shelef, M., Iwakoshi, N.N., Lee, A.H., Qian, S.B., Zhao, H., Yu, X., Yang, L., Tan, B.K., Rosenwald, A., et al. (2004). XBP1, downstream of Blimp-1, expands the secretory apparatus and other organelles, and increases protein synthesis in plasma cell differentiation. Immunity 21, 81–93.

Shinnakasu, R., Inoue, T., Kometani, K., Moriyama, S., Adachi, Y., Nakayama, M., Takahashi, Y., Fukuyama, H., Okada, T., and Kurosaki, T. (2016). Regulated selection of germinal-center cells into the memory B cell compartment. Nat Immunol 17, 861–869.

Stone, S.L., Peel, J.N., Scharer, C.D., Risley, C.A., Chisolm, D.A., Schultz, M.D., Yu, B., Ballesteros-Tato, A., Wojciechowski, W., Mousseau, B., et al. (2019). T-bet Transcription Factor Promotes Antibody-Secreting Cell Differentiation by Limiting the Inflammatory Effects of IFN-gamma on B Cells. Immunity.

Subramanian, A., Tamayo, P., Mootha, V.K., Mukherjee, S., Ebert, B.L., Gillette, M.A., Paulovich, A., Pomeroy, S.L., Golub, T.R., Lander, E.S., and Mesirov, J.P. (2005). Gene set enrichment analysis: a knowledge-based approach for interpreting genome-wide expression profiles. Proc Natl Acad Sci U S A 102, 15545–15550.

Takahashi, Y., and Kelsoe, G. (2017). Role of germinal centers for the induction of broadly-reactive memory B cells. Curr Opin Immunol 45, 119–125.

Taubenheim, N., Tarlinton, D.M., Crawford, S., Corcoran, L.M., Hodgkin, P.D., and Nutt, S.L. (2012). High rate of antibody secretion is not integral to plasma cell differentiation as revealed by XBP-1 deficiency. J Immunol 189, 3328–3338.

Tibshirani R, W.G., Hastie T (2001). Estimating the number of clusters in a data set via the gap statistic. Journal of the Royal statistical society: series B (Methodological) 63, 411–423.

Tipton, C.M., Fucile, C.F., Darce, J., Chida, A., Ichikawa, T., Gregoretti, I., Schieferl, S., Hom, J., Jenks, S., Feldman, R.J., et al. (2015). Diversity, cellular origin and autoreactivity of antibody-secreting cell population expansions in acute systemic lupus erythematosus. Nat Immunol 16, 755–765.

Tsukumo, S., Unno, M., Muto, A., Takeuchi, A., Kometani, K., Kurosaki, T., Igarashi, K., and Saito, T. (2013). Bach2 maintains T cells in a naive state by suppressing effector memory-related genes. Proc Natl Acad Sci U S A 110, 10735–10740.

Turner, J.S., Zhou, J.Q., Han, J., Schmitz, A.J., Rizk, A.A., Alsoussi, W.B., Lei, T., Amor, M., McIntire, K.M., Meade, P., et al. (2020). Human germinal centres engage memory and naive B cells after influenza vaccination. Nature 586, 127–132.

van Anken, E., Romijn, E.P., Maggioni, C., Mezghrani, A., Sitia, R., Braakman, I., and Heck, A.J. (2003). Sequential waves of functionally related proteins are expressed when B cells prepare for antibody secretion. Immunity 18, 243–253.

Wang, S., Wang, J., Kumar, V., Karnell, J.L., Naiman, B., Gross, P.S., Rahman, S., Zerrouki, K., Hanna, R., Morehouse, C., et al. (2018). IL-21 drives expansion and plasma cell differentiation of autoreactive CD11c(hi)T- bet(+) B cells in SLE. Nat Commun 9, 1758.

Woodland, D.L., and Kohlmeier, J.E. (2009). Migration, maintenance and recall of memory T cells in peripheral tissues. Nat Rev Immunol 9, 153–161.

Wrammert, J., Smith, K., Miller, J., Langley, W.A., Kokko, K., Larsen, C., Zheng, N.Y., Mays, I., Garman, L., Helms, C., et al. (2008). Rapid cloning of high-affinity human monoclonal antibodies against influenza virus. Nature 453, 667–671.

Xu, H., Chaudhri, V.K., Wu, Z., Biliouris, K., Dienger-Stambaugh, K., Rochman, Y., and Singh, H. (2015). Regulation of bifurcating B cell trajectories by mutual antagonism between transcription factors IRF4 and IRF8. Nat Immunol 16, 1274–1281.

Xu, W., Zhao, X., Wang, X., Feng, H., Gou, M., Jin, W., Wang, X., Liu, X., and Dong, C. (2019). The Transcription Factor Tox2 Drives T Follicular Helper Cell Development via Regulating Chromatin Accessibility. Immunity 51, 826–839 e825.

Y, B.Y.a.H. (1995). Controlling the fase discovery rate: a practical and powerful approach to multiple testing. . Journal of the Royal statistical society: series B (Methodological) 57, 289–300.

Yao, C., Lou, G., Sun, H.W., Zhu, Z., Sun, Y., Chen, Z., Chauss, D., Moseman, E.A., Cheng, J., D’Antonio, M.A., et al. (2021). BACH2 enforces the transcriptional and epigenetic programs of stem-like CD8(+) T cells. Nat Immunol.

Yu, B., Zhang, K., Milner, J.J., Toma, C., Chen, R., Scott-Browne, J.P., Pereira, R.M., Crotty, S., Chang, J.T., Pipkin, M.E., et al. (2017). Epigenetic landscapes reveal transcription factors that regulate CD8(+) T cell differentiation. Nat Immunol 18, 573–582.

Zhang, Y., Liu, T., Meyer, C.A., Eeckhoute, J., Johnson, D.S., Bernstein, B.E., Nusbaum, C., Myers, R.M., Brown, M., Li, W., and Liu, X.S. (2008). Model-based analysis of ChIP-Seq (MACS). Genome Biol 9, R137.

Zhou, X., and Xue, H.H. (2012). Cutting edge: generation of memory precursors and functional memory CD8+ T cells depends on T cell factor-1 and lymphoid enhancer-binding factor-1. J Immunol 189, 2722–2726.

Zhou, X., Yu, S., Zhao, D.M., Harty, J.T., Badovinac, V.P., and Xue, H.H. (2010). Differentiation and persistence of memory CD8(+) T cells depend on T cell factor 1. Immunity 33, 229–240.

Zumaquero, E., Stone, S.L., Scharer, C.D., Jenks, S.A., Nellore, A., Mousseau, B., Rosal-Vela, A., Botta, D., Bradley, J.E., Wojciechowski, W., et al. (2019). IFNgamma induces epigenetic programming of human T-bet(hi) B cells and promotesTLR7/8 and IL-21 induced differentiation. Elife 8.

